# Precise timing is ubiquitous, consistent and coordinated across a comprehensive, spike-resolved flight motor program

**DOI:** 10.1101/602961

**Authors:** Joy Putney, Rachel Conn, Simon Sponberg

## Abstract

Sequences of action potentials, or spikes, carry information in the number of spikes and their timing. Spike timing codes are critical in many sensory systems, but there is now growing evidence that millisecond-scale changes in timing also carry information in motor brain regions, descending decision-making circuits, and individual motor units. Across all the many signals that control a behavior how ubiquitous, consistent, and coordinated are spike timing codes? Assessing these open questions ideally involves recording across the whole motor program with spike-level resolution. To do this, we took advantage of the relatively few motor units controlling the wings of a hawk moth, *Manduca sexta*. We simultaneously recorded nearly every action potential from all major wing muscles and the resulting forces in tethered flight. We found that timing encodes more information about turning behavior than spike count in every motor unit, even though there is sufficient variation in count alone. Flight muscles vary broadly in function as well as in the number and timing of spikes. Nonetheless, each muscle with multiple spikes consistently blends spike timing and count information in a 3:1 ratio. Coding strategies are consistent. Finally, we assess the coordination of muscles using pairwise redundancy measured through interaction information. Surprisingly, not only are all muscle pairs coordinated, but all coordination is accomplished almost exclusively through spike timing, not spike count. Spike timing codes are ubiquitous, consistent, and essential for coordination.

**Significance Statement:** Brains can encode precise sensory stimuli and specific motor systems also appear to be precise, but how important are millisecond changes in timing of neural spikes across the whole motor program for a behavior? We record every spike that the hawk moth’s nervous system sends to its wing muscles. We show that all muscles convey the majority of their information in spike timing. The number of spikes does play a role, but not in a coordinated way across muscles. Instead, all coordination is done using in the millisecond timing of in spikes. The importance and prevalence of timing across the motor program pose new questions for how nervous systems create precise, coordinated motor commands.

---

Neurons convey information not only through the number of spikes, but also their timing (1–4). In sensory systems, both changes in the number of spikes over time and precise, millisecond-level shifts in sequences of spikes are well established as essential encoding mechanisms for proprioception (5), audition (6), vision (1, 7–9), touch (10), and other modalities (7, 11, 12). Spike timing codes have been shown to be of particular importance in sensory systems (1, 7, 8), and patterns of multiple spikes can convey more information about a stimulus than the sum of the individual timings (13). In vertebrate motor systems, rate codes, where muscle force is proportional to the firing rate of the motor neuron, are thought to predominate, in part due to recruitment principles of many motor units and the presumed low-pass nature of muscles (14–17). Although vertebrate muscle force may be modulated by spike rate under isometric conditions (15), precisely timed patterns of spikes affect the output force of muscle (18). Similarly, in invertebrates, rate codes can adjust force development in muscles, but the absolute number of spikes (spike count code) also matters (19, 20). The onset time a single spike or burst is also known to play a functional role for the control of invertebrate muscle (21–24).

Recent evidence in invertebrates and vertebrates shows that spike timing codes may be under-appreciated for controlling motor behaviors at least in specific muscles or motor circuits (4). spike timing codes in which information is encoded in the precise timing patterns of neural or muscular action potentials have an even higher capacity to code for the output of muscles than rate or count (4, 13, 18). Such codes are found in a songbird cortical area for vocalization (25) and in mouse cerebellum for task error correction (26). Correlational, causal, and mechanistic studies show that millisecond-level changes in timing of spikes in motor neurons can manifest profound changes in force production (27) and even behavior selection (28). Causal evidence in support of spike timing codes is present in fast behaviors like invertebrate flight (27), but also in relatively slow behaviors like breathing in birds (18). However, evidence for the importance of spike timing codes in motor systems has been limited to only a few of the motor signals that typically control movement. Whether such timing codes are utilized broadly across a complete motor program for behavior is unknown as is their role in coordinating multiple motor units. Despite growing appreciation of the potential for motor timing codes, we have not yet established the ubiquity, consistency and coordination of spiking timing across the motor signals that compose a behavior. This poses three hypotheses.

First, timing codes may be restricted to only a few motor signals that control behavior. For example, recordings of muscles in locusts, hawk moths, and fruit flies have shown that spike timing and count variation are prevalent in specific motor units (21, 22, 29). Alternatively, timing codes may be ubiquitous–widespread across the entire motor program and present in all muscles controlling a behavior.

Regardless of the prevalence of timing codes, motor neurons within the population may exhibit specialized encoding strategies, varying the amount of information transmitted through spike timing or spike count depending on the function of the muscles they innervate. For example, *Drosophila* use combinations of functionally distinct phasic and tonic motor units to control flight (23). Additionally, evidence in some sensory systems show separate classes of neurons use either spike rate or spike timing to convey information (30). Alternatively, the entire motor program may be consistent in its use of spike timing for encoding.

Finally, coordination of multiple motor signals is typically assessed through covariation in firing rates. For example, motor coordination patterns across muscles (*e.g.* muscle synergies (31)) and population recordings of M1 neurons in motor cortex (32) all consider how populations of units encode movement through spike rate. Alternatively, spike timing codes may play a role in the coordination of muscles in motor systems. Resolving these hypotheses about the role of spike timing in motor control is challenging because they consider encoding strategies across an entire motor program. It is therefore necessary to record from a spike-resolved, comprehensive set of signals that control a behavior simultaneously in a consistent behavioral context.

Recording such a comprehensive motor program is difficult due to the requirements of completeness, sufficient temporal resolution, and sampling rich variation in a naturalistic behavior. Obtaining a nearly complete motor program is more tractable in the peripheral nervous system than in central regions because of smaller neuronal population sizes. While many muscles or motor units have been simultaneously recorded using electromyography (EMG) in frogs (36), cats (31), and humans (37) and using calcium imaging in the wing steering muscles of fruit flies (23), these sets of neural signals are not spike-resolved. Large flying insects are feasible organisms in which to record a spike-resolved, comprehensive motor program because all muscles actuating the wings are in the thorax and there are relatively few muscles compared to many segmented limbs. Moreover flight muscles frequently function as single motor units because they are generally innervated by one or very few fast-type motor neurons with a 1:1 relationship between muscle and neural potentials (38, 39). A large number of spike-resolved motor units has been simultaneously recorded in locusts (40) and a smaller number simultaneously in flies and moths (41–43), although explicit analysis of encoding in count and timing has not been done in these systems. Invertebrate muscles have distinct count (number of spikes) and rate codes that do not have interchangeable effects on muscle force (19), but both of these are distinct from spike timing coding. Faster invertebrate muscles fire fewer times per cycle but can still show rate coding during and across wing strokes (29).

We take advantage of these features to capture a spike-resolved, comprehensive motor program in a hawk moth, *Manduca sexta*, and investigate the importance of spike timings in a nearly complete population code for movement. We examine how turning torque in every wing stroke is encoded by spike count (the number of spikes per wing stroke) and spike timing (the precise timing patterns of all spikes within each wing stroke) for each of the 10 muscles most important for controlling the wings (SI Appendix) (21, 44–46). This nearly complete motor program enables us to address three questions of ubiquity, consistency, and coordination in timing and count codes across this motor system.

## Results

### Temporal information is ubiquitous in the motor program

We recorded a comprehensive motor program with spike-level resolution across all the primary muscles actuating the wings in a hawk moth (*Manduca sexta*, N = 7) (Fig. 1*A*). The hawk moth musculature has been examined in detail anatomically and through *in vivo* and *in vitro* recordings (summarized in SI Appendix). Based on this rich literature we identified five bilateral pairs of muscles that have important roles in controlling the wings during flight (Fig. S1). We recorded EMG signals from these muscles while moths visually tracked a robotic flower in tethered, smooth pursuit flight (27, 47). We simultaneously recorded within wing stroke yaw torque using a custom calibrated force-torque transducer (ATI Nano17Ti) (Fig. 1*B-D*). We segmented the EMG and torque data into wing strokes. We defined the onset time of each wing stroke as the zero phase crossing of the Hilbert transform of the moth’s force in the z-direction. The Hilbert transform estimates the phase of a periodic signal; here the zero phase crossing roughly corresponded to the peak downward force produced during each wing stroke. We treated each wing stroke as an independent sample of the muscle spikes and the yaw torque.

**Fig. 1.**
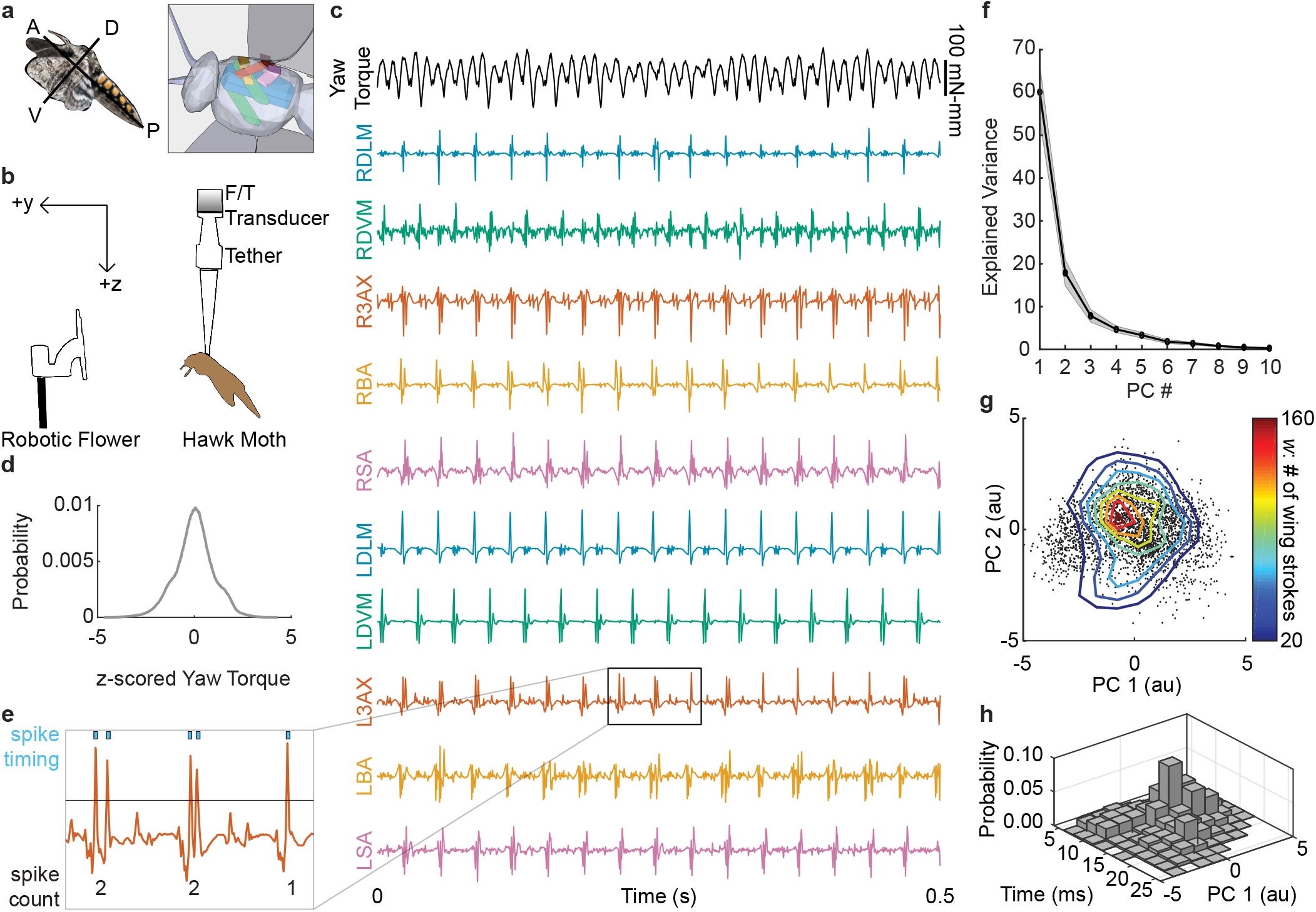
EMGs from 10 flight muscles and simultaneous yaw torque. (*A*) A hawk moth, *Manduca sexta*, in flight with a simplified 3D sketch of the 5 bilateral pairs of muscles from a ventrolateral view: dorsolongitudinal, DLM (blue); dorsoventral, DVM (green); 3rd axillary, 3AX (orange); basalar, BA (yellow); subalar, SA (purple). Muscles on the left and right sides of the animal are distinguished with an L or an R throughout the text (ex. L3AX). (*B*) Hawk moths were presented a robotic flower oscillating with a 1 Hz sinusoidal trajectory while tethered to a custom six-axis F/T transducer (N = 7 moths; 999-2,954 wing strokes per moth; average per moth = 1,950 wing strokes). (*C*) EMG and yaw torque (black) from 0.5 seconds of flight. (*D*) A population-level histogram of the raw yaw torque during shortened wing strokes z-scored in each individual moth. (*E*) Spike sorting was accomplished using threshold crossing (e.g. black line) in Offline Sorter (Plexon). Spike count is the number of spikes in each wing stroke, and spike timing is the precise spike time relative to the start of each wing stroke. (*F*) The first two principal components (PCs) of the yaw torque waveforms captured most of the variance (mean, in black; *±* S.E.M., in gray; N = 7 moths). (*G*) Projection of yaw torque onto the first two PCs for each wing stroke from a moth (*w* = 2,739 wing strokes) in PC space (arbitrary units, *au*). The joint histogram of the distribution is represented in a 10 × 10 grid between −5 and 5 using isoclines from the contour function in MATLAB (MathWorks). (*H*) Joint histogram of the scores of PC1 and the timing of the LDLM spike in wing strokes in an example moth.

For the EMG data, we specified a time window relative to the onset of the wing stroke separately for each muscle to encompass the entire burst of spikes in all wing strokes. We computed the spike count (number of spikes per wing stroke) or the spike timing (precise spike times relative to the start of each wing stroke) for the 10 muscles (Fig. 1*E*). Because the wing stroke period varied a small amount (mean *±* s.d.: 45.3 *±* 3.8 ms across all moths), we shortened the yaw torque signal to the length of the shortest wing stroke for each moth. We also repeated our analyses with a phase code (timing normalized to wingstroke period) and obtained similar results. We then found a lower dimensional representation of the yaw torque using principal components analysis (PCA). The first two PCs explained most of the variance (78.0 ± 10.6%) in yaw torque (Fig. 1*F*). The oscillating visual stimulus elicited variation in the moths’ motor output and spiking activity (Fig. 1*G-H*). The scores of the two PCs correspond with left and right turns in the extreme deciles of movement and captured the main features of the torque timeseries (Fig. 2).

**Fig. 2.**
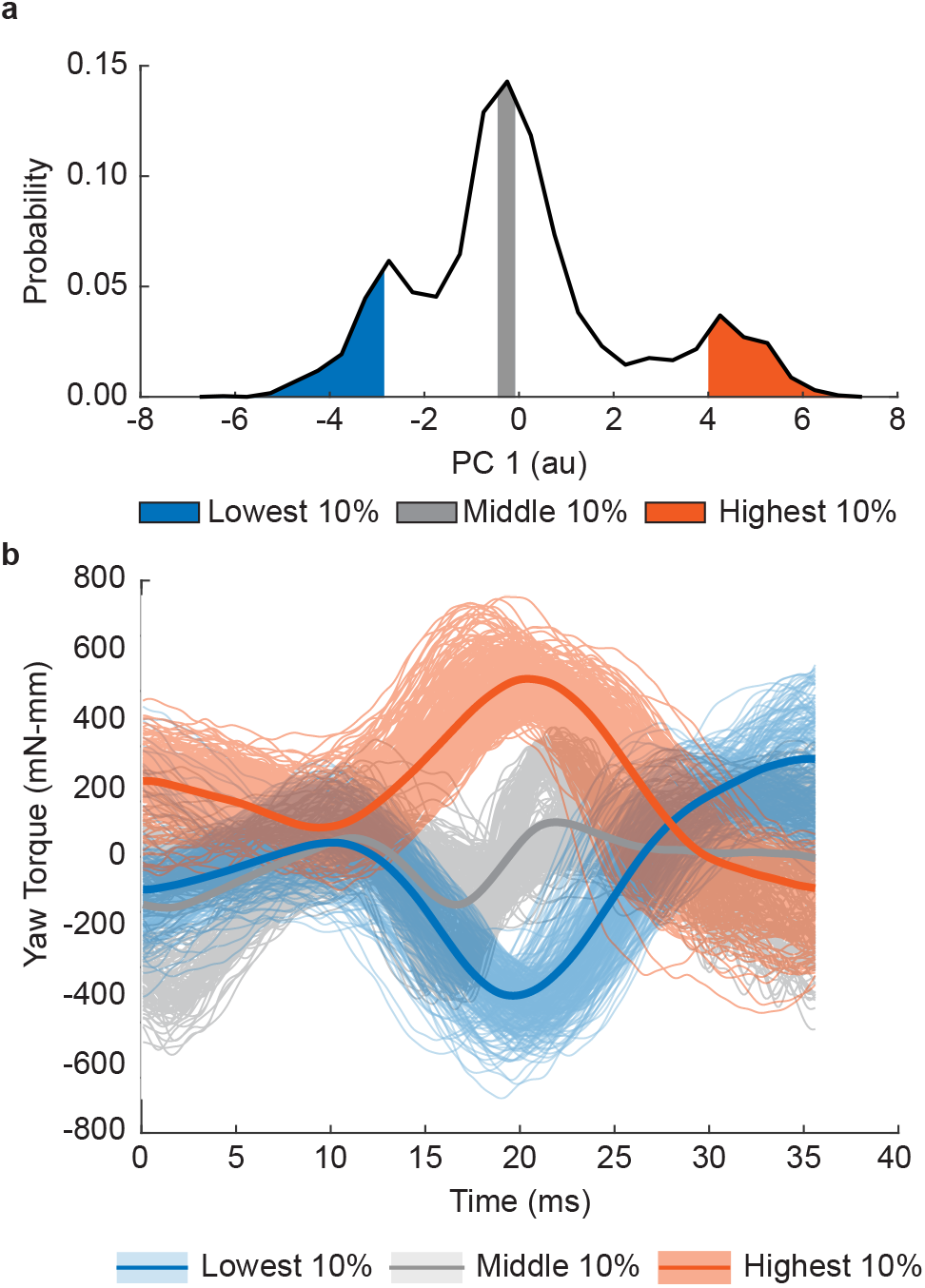
Reconstructions using the first 2 PCs of yaw torque capture the main features of the raw torque waveforms. (*A*) Histogram of the scores of PC1 in an individual moth. The lowest decile (0-10%), the middle decile (45-55%), and highest decile (90-100%) are shaded in blue, grey, and orange, respectively. (*B*) The average reconstruction of wing strokes (solid lines) from the lowest (blue), middle (grey), and highest (orange) deciles using the data set mean and the projections of scores onto the first 2 PCs, along with the raw torque waveforms (translucent lines) in each of these deciles. Deciles vary both in mean torque and the within wing stroke torque waveforms consistent with early methods (33).

Both the spike count and the timing of spikes within the wing stroke show modulation along with the motor output (Fig. 3*A*). To test the contribution spike timing encoding in individual muscles, we estimated the mutual information between muscle activity and yaw torque using the Kraskov *k*- nearest neighbors method, which is data efficient and useful for experiments where sampling is finite and measured variables are continuous (34, 35). Unlike a direct method estimator, this method estimates mutual information between two variables (*X* and *Y*) using *k*-nearest neighbor Euclidean distances between each sample (wing stroke) and its *k*th nearest neighbor in the space spanning the two variables of interest (for us, a representation of spiking activity and torque). The joint probability distribution of the distances and the number of samples within a neighborhood defined by these distances is used to estimate the joint entropy *H*(*X, Y*) and the mutual information *I*(*X, Y*).

**Fig. 3.**
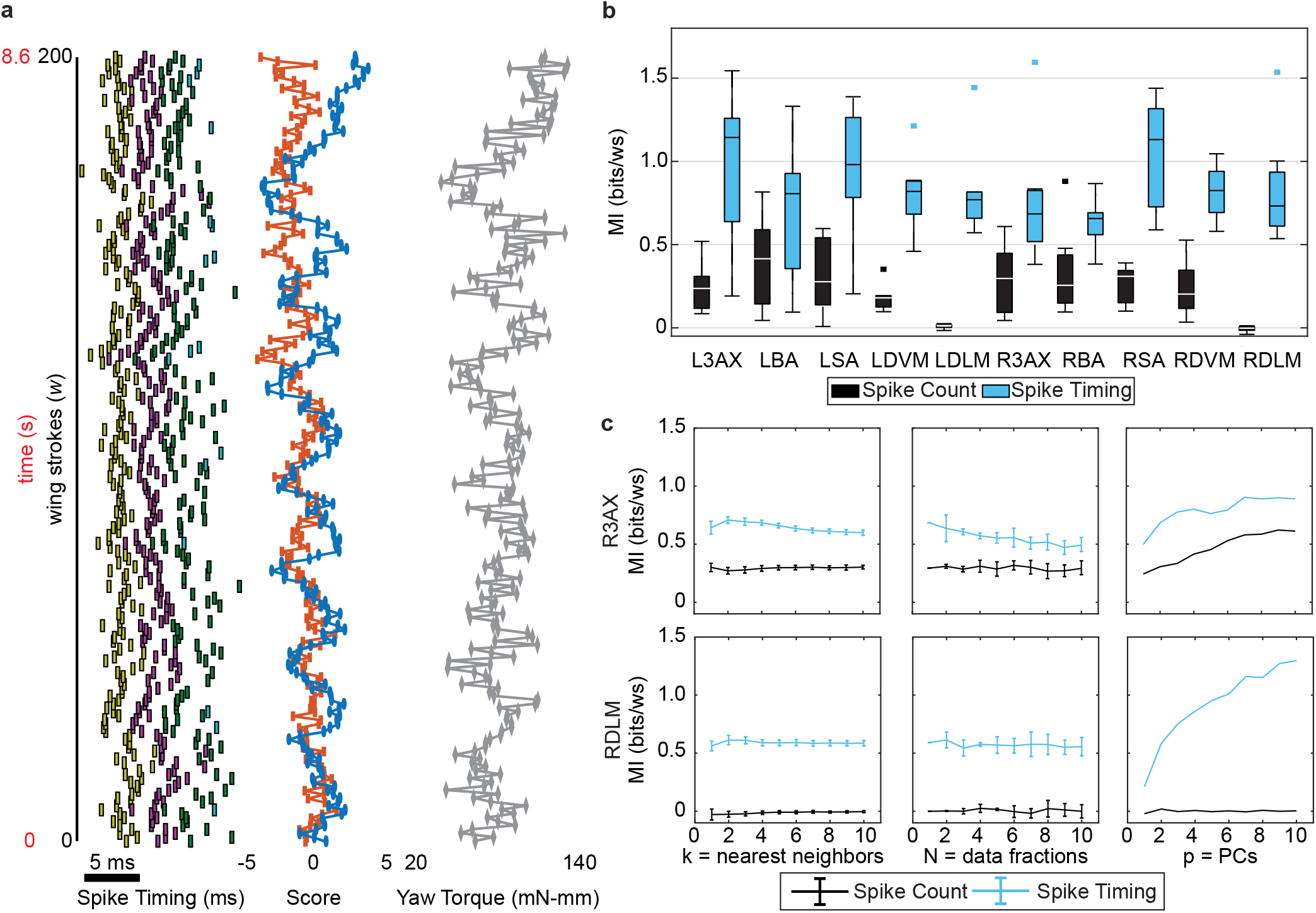
Mutual information between spike count or spike timing and yaw torque. (*A*) Timing of spikes in the L3AX, the scores of the first 2 PCs, and the wing stroke average yaw torque show variability corresponding with the 1 Hz visual stimulus (200 wing strokes from a moth are shown). The rasters are the 1st (yellow), 2nd (purple), 3rd (green), and 4th (light blue) spikes within each wing stroke shown alongside the 1st (blue) and 2nd (red) yaw torque PC scores and the raw yaw torque. (*B*) MI estimates for spike count (black) and spike timing (blue) with yaw torque across individuals (N = 7). Box plots report the median as the center line in the box, which marks the 25th and 75th percentiles. Whiskers are the range of all points that are not considered outliers (square points). Spike count MI is less than spike timing MI (two-way ANOVA comparing timing vs. count for all muscles: count vs. timing, p *<* 10^*−*10^; muscle ID, p = 0.26; interaction, p = 0.09). Spike timing MI is significantly greater than spike count MI in most paired comparisons within muscles (paired t-tests: p *<* 0.02 for all muscles except the LBA, p = 0.09, and RBA, p = 0.05. Wilcoxon signed rank tests: p *<* 0.02 for all muscles except the LBA, p = 0.11, and RBA, p = 0.08). (*C*) MI estimates (mean *±* S.D.) for the number of nearest neighbors *k* = 1-10, data fractions *N* = 1-10, and PCs included *p* = 1-10 from the RDLM and R3AX muscles of one moth (34, 35).

This information theoretic approach enables us to consider the importance of spike timing without assuming what features of the spike train are relevant or a linear relationship between spiking and motor output (48). It also enables separate of spike count mutual information (MI) from spike timing MI by conditioning spike timing on spike count (18):

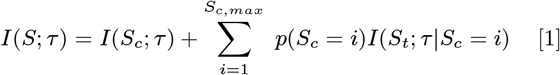

where *S* is the combined representation of spike count and precise timing within the wingstroke of each spike, and *τ* is the 2 variable representation of motor output taken as the scores of the two yaw torque PCs. *S*_*c*_ is the spike count for each wing stroke taking discrete states, *i*, from 1 to *S*_*c,max*_. *S*_*t*_ is a vector of spike timing variables conditioned upon *S*_*c*_ such that for each spike count it has the same length, *i*. The first term is the mutual information between torque and count. The second term is the mutual information between torque and timing once the information in count is accounted for.

For all 10 muscles, spike timing MI is higher than spike count MI (Fig. 3*B*). In all muscles both spike count MI and spike timing MI are non-zero, except for the DLM, which only spikes once per wing stroke during flight (range of mean spike count MI across 10 muscles = 0.0 - 0.4 bits/wing stroke (ws); spike timing MI = 0.6 - 1 bits/ws). All muscles in the motor program that vary the number of spikes present in each wing stroke use mixed encoding, a combination of spike timing and spike count, to inform the torque. The error estimates (see Methods) of the MIs were small compared to the total MI (Table S1, mean spike count and timing MI error *<* 0.04 bits/ws across all muscles). Our MI estimates are stable across varying values of *k*, the number of nearest neighbors, and the number of data fractions (Fig. 3*C*, S2, S3). In the spike timing MI estimations, 90% of estimations from halved data sets deviated by less than 10% from the full data set estimate.

Temporal encoding is ubiquitous across the entire flight motor program, present in every muscle, and is utilized more than count encoding (Fig. 3*B*). Each motor unit encodes almost an order of magnitude more information per period about yaw torque in precise spike timings (0.8 bits/ws on average for all muscles) compared to other systems, like a cortical vocal area (between 0.1-0.3 bits/syllable) (25) and breathing muscles (between 0.05-0.2 bits/breath cycle) of song birds (18). The moth’s 10 motor units collectively code for flight using on the order of 1 bit per wing stroke each.

### Encoding strategy is consistent across functionally diverse muscles

Muscles in the hawk moth motor program have diverse biomechanical functions. For example, the main indirect downstroke muscle (dorsolongitudinal muscle, DLM), acts by pulling on the exoskeleton to contract the thorax, propagating mechanical strain to the wing hinge and causing the wings to depress (21). In contrast to the DLM, the third axillary muscle (3AX) directly attaches to the wing hinge at the third axillary sclerite, which articulates the anal vein, the most posterior vein of the forewing (46, 51). Muscles also have variable spiking activity. Different muscles have different probability distributions of spike count per wing stroke (i.e. spike rate) and spike timing during the wing strokes (Fig. 4*A-B*).

**Fig. 4.**
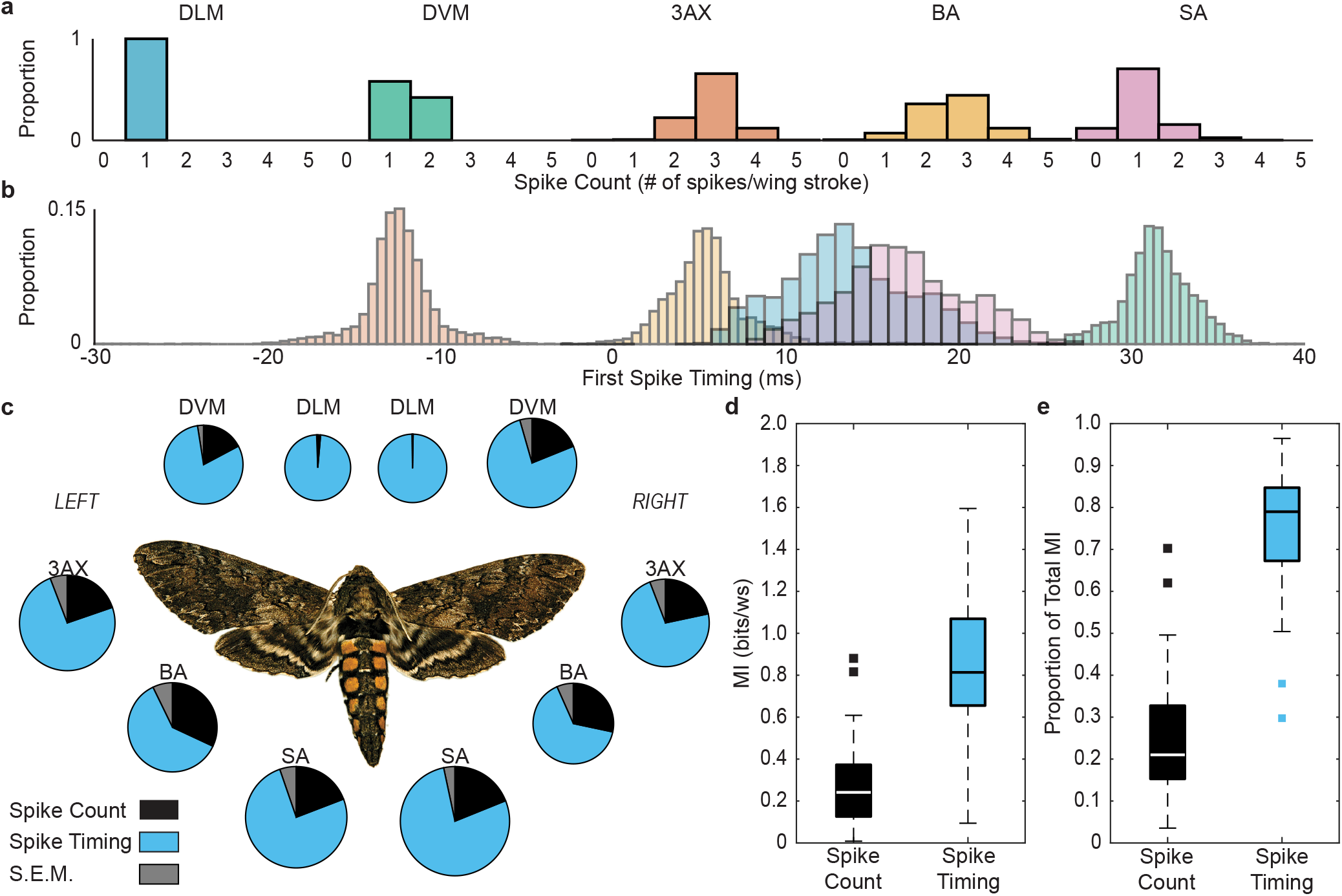
Consistency of magnitude and proportion of spike timing MI and spike count MI in all 10 muscles. The 5 muscle types we recorded have different probability distributions of spike count conditions (*A*) and the first spike timing (*B*) (data shown for one moth). Some bursts begin before the wing stroke and continue into the wing stroke; these were reported as negative values (t = 0 corresponds to the start of the wing stroke). (*C*) Mean spike count and spike timing MI estimates for all 10 muscles across individuals (N = 7). Pie size indicates the magnitude of total MI, and the slices indicate the proportion that is spike count MI (black) and spike timing MI (blue), as well as the S.E.M. these proportions (gray). No significant difference was found in the magnitude of spike count MI of all muscles excluding the DLM (one-way ANOVA: p = 0.66; Kruskal-Wallis test: p = 0.90) or spike timing MI of all muscles (one-way ANOVA: p = 0.54; Kruskal-Wallis test: p = 0.39). No significant difference was found in the proportion of spike timing MI to total MI in all muscles excluding the DLM (one-way ANOVA: p = 0.31; Kruskal-Wallis test: p = 0.54). (*D-E*) The magnitude or proportion of spike count MI (black) and spike timing MI (blue), respectively, across 8 muscles (DLM excluded) and 7 individuals. Boxplots display data as previously described in Fig. 3*B*.

Despite their diverse properties, the 10 muscles in the motor program of the hawk moth are consistent in the magnitude and proportion of timing information used to encode yaw torque (Fig. 4*C*). No muscle conveys significantly different spike timing MI. Additionally, all muscles that spike more than once per wing stroke carry similar amounts of spike count MI. As a result, there is a consistent 3:1 ratio of spike timing MI to spike count MI for all muscles that spike more than once per wing stroke (Fig. 4*C-E*: mean ± 95% C.I. of the mean of the ratio of spike timing MI to total MI for all muscles excluding DLM = 0.75 *±* 0.02).

Our conclusions were robust if we reduced the representation of the yaw torque to the scores of just the first PC (Fig. S4*A*) or the average torque during a wing stroke (Fig. S4*B*). Increasing dimensional to 3 PCs (Fig. S5) destabilizes estimates of information in some muscles due to data limits, but our conclusions nonetheless remain consistent.

Neurons in some sensory systems may use distinct strategies to encode particular types of information (30). However, this is not the case in the hawk moth motor program. Even though each muscle has a different probability distribution of spike count and spike timing (Fig. 4*A-B*), each muscle shares a comparable amount of MI with the moth’s torque. The different probability distributions may indicate that different muscles have varying amounts of total entropy (bandwidth) while transmitting the same amount of information. Alternatively, different muscle types may have comparable total entropies, but encode torque with varying precision.

### Coordination is achieved through timing, not count

Because timing is ubiquitous across all muscles and encoding strategies are consistent, we next investigated the role of spike timings in the coordination of multiple muscles. To do this, we first estimated the pairwise MI between the spiking activity of two muscles and the yaw torque:

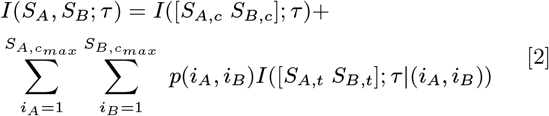

*I*(*S*_*A*_, *S*_*B*_; *τ*) is the pairwise MI, or the mutual information between the torque and the joint spiking activity of two muscles, *S*_*A*_ and *S*_*B*_. As before (Eq. 1), the first term is the pairwise spike count MI, and the second term is the pairwise spike timing MI. The estimates are weighted by the joint probability *p*(*i*_*A*_, *i*_*B*_) of each possible pairwise spike count condition. As in the individual MI estimations, we used a value of *k* = 4 (Fig. S7).

Then, we estimated the interaction information (*II*) between two muscles (49, 50):

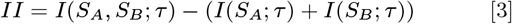

A positive *II* indicates net synergistic information, or that the muscles together reduce the entropy of the motor output more than the sum of their individual contributions. A negative *II* indicates that information is net redundant between the two muscles, or that there is coordination in the information content between the two muscles.

All pairwise combinations of muscles in the motor program have non-zero, negative *II* values (Eq. (3)), which are net redundant interactions (Fig. 5*A*). We separated the contribution of count from the contribution of timing information to *II* as *II*_*count*_ and *II*_*timing*_ (Eq. S1-S2 in SI Methods) and found that nearly all redundant information between muscles is encoded in spike timing (Fig. 5*B-C*, S6). Mean spike count *II* is −0.023 ± 0.006 bits/ws, while mean spike timing *II* is −0.56 ± 0.04 bits/ws (± 95% C.I. of the mean). Spike timing, not count accomplishes essentially all of the coordination between muscles in the motor program.

**Fig. 5.**
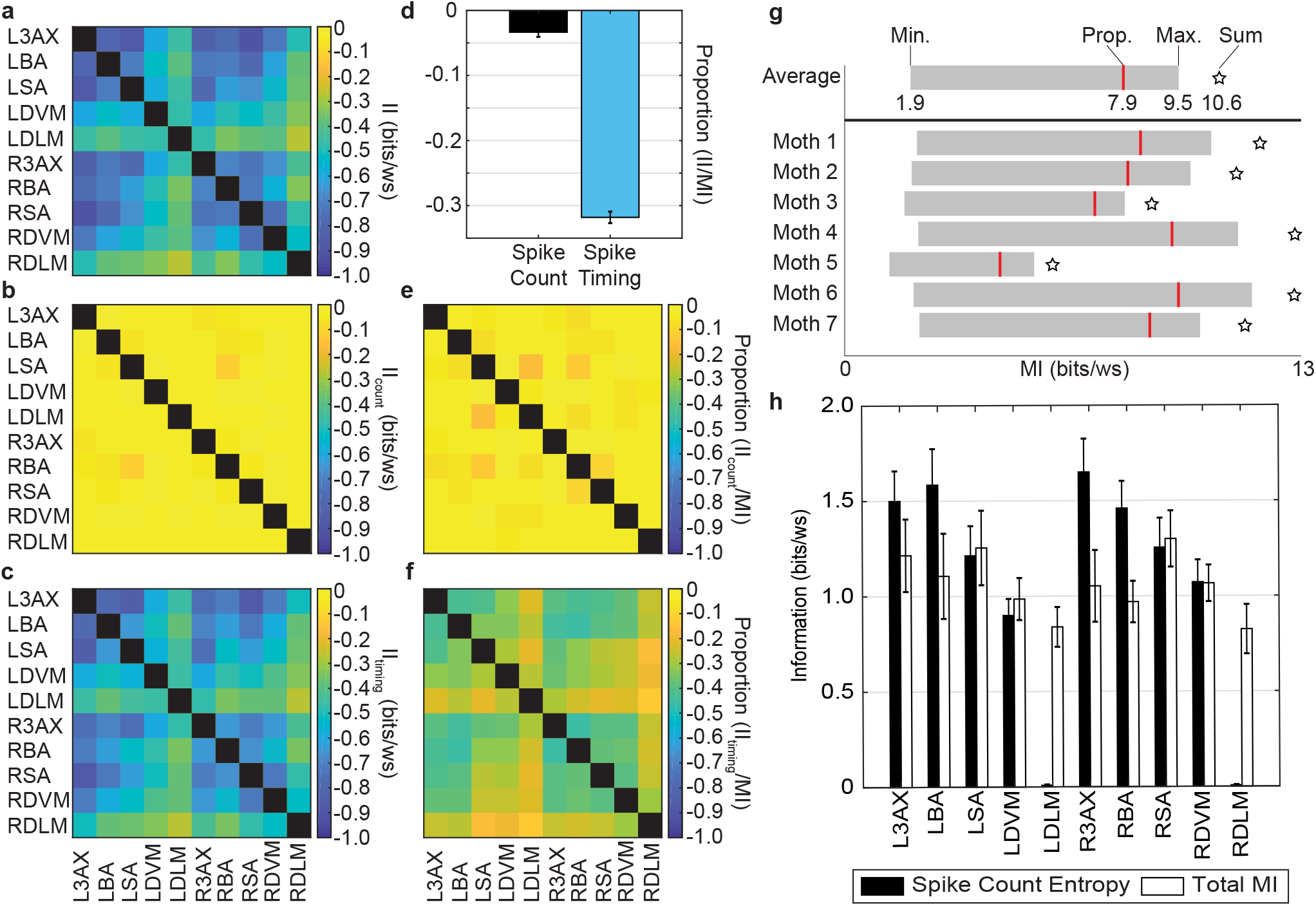
Interaction information in pairwise combinations of muscles and the range of total motor program MI values possible. (*A*) We calculated total interaction information (*II*) (Eq. (3)) (49, 50) as a measure that compares the estimates of pairwise MI (Eq. (2)) and individual muscle MI (Eq. (1)) for all pairwise combinations of muscles (mean for N = 7 moths). All values of *II* are negative, indicating net redundant interactions or overlapping information content. Comparisons of muscles to themselves are excluded. (*B-C*) Spike count interaction information (*II*_*count*_) or spike timing interaction information (*II*_*timing*_), respectively, across all pairwise combinations of muscles (Eq. S1 and Eq. S2 in SI Methods, mean for N = 7). (*D*) Proportion of *II* to the sum of individual muscle MIs for spike count and timing terms of Eq. S1-S2 (mean *±* S.E.M., all muscle pairs excluding DLMs, n = 56). (*E-F*) The proportion of *II*_*count*_ or *II*_*timing*_ to the sum of the individual spike count or timing MIs, respectively (mean for N = 7 individuals). (*G*) Estimates of lowest and highest values of total motor program MI (gray box), proportional estimate of motor program MI (red line), and sum of individual muscle MIs (star) for each moth and the population average. (*H*) Mean *±* S.E.M. of the spike count entropy (Eq. S3 in SI Methods) and the total MI (N = 7).

It is possible spike timing is more important for coordination than count simply because spike timing encodes more information overall. To test this we scaled the spike count and spike timing interaction information according to the total magnitude of spike count and spike timing mutual information. Overall, 31.8 ± 0.9% of spike timing MI and 3.4 ± 0.9% of spike count MI in individual muscles is shared in pairwise interactions (Fig. 5*D*). Even considering the smaller magnitude of spike count MI in individual muscles, spike count encodes almost no coordinated information (Fig. 5*E-F*). Count encoding of each muscle is independent of other muscles in the motor program.

### The motor program utilizes less than 10 bits/wing stroke

Coordination between muscles and limited amounts of information in each muscle suggest that the motor program operates with no more than 10 bits of information per wing stroke. We created maximum, minimum, and intermediate (balanced) estimates of the total information content of the motor program, using several methods to account for the redundant information in pairwise combinations (Eq. S4-S5 in SI Methods).

The comprehensive flight motor program uses a MI rate between 1.85 bits/ws to 9.47 bits/ws, with an intermediate estimate of 7.89 bits/ws (Fig. 5*G*). Since the average wing stroke length used in these calculations was 0.04 s, this corresponds to an information rate between 46.2 bits/s to 237 bits/s. Lacking other comprehensive motor program recordings it is difficult to compare information rate across motor systems. However, hawk moth flight is accomplished with a small information rate compared to those in sensory systems. While individual sensory neurons have comparable information rate (6-13 bits/s in RGCs (52) and 1-10 bits/s in olfactory receptors (53)), these systems have orders of magnitude more receptors, so the maximum information rate is orders of magnitude higher overall (875,000 bits/s in the guinea pig retina (52)).

This information rate allows the moth to specify a large number of possible motor outputs. To estimate this range of motor output, we determined how many states in the empirical torque probability distribution could be encoded by the total motor program using the direct method (Eq. S6 in SI Methods). Given the intermediate estimate between the upper and lower values, the motor program MI can specify 483 ± 109 states of yaw torque (N = 7 individuals) for each wing stroke. We also estimated the entropy in spike count using the direct method (Eq. S3 in SI Methods). Excluding the DLM, the count entropy in each muscle was as least as large as the total MI (Fig. 5*H*). With noiseless transmission, the motor program could be encoded strictly in count.

## Discussion

By investigating a comprehensive, spike-resolved motor program, we show that spike timing encoding is not a feature of just specialized motor units, but a ubiquitous control strategy that is consistently used for activation and coordination of muscles. There are few, if any, differences in magnitudes and proportions of spike timing and spike count encoding between the various muscles controlling the wings (Fig. 3*B*, 4), despite their different modes of actuation and functional diversity (21). All muscles encode information about yaw torque in both precise spike timing and spike count (Fig. 4*C-E*). Spike count is significant in every muscle with the exception of the DLMs which only spike once per wing stroke during flight. However, when it comes to coordination between pairwise combinations of muscles, timing is almost everything.

The moth motor program has individual muscles acting as mixed spike timing and spike count encoders. *In situ* preparations of a wing elevator muscle in a locust, *Schistocerca nitens*, showed that changing either the spike timing or the number of spikes altered power output (54). Steering muscles, like the basalar muscle in the blowfly *Calliphora vicina*, can act by dissipating energy rather than doing positive work and the timing of activation can modulate power (55). In this species, timing in the basalar muscle and coordination between pairs of activated muscles have been shown to effect wing kinematics and total body force (42, 43). Most of the mechanistic studies to date have examined how activation signals of a subset of muscles affect muscle force or body movements, but comprehensive stimulation investigating the effects of coordinated control mechanisms across muscles will be needed to understand functional implications. From our results, we now know any studies of the moth’s complete motor program must examine spike timing.

Spike timing can still matter in vertebrate muscle because of non-linearity in force development and biomechanics (4). By shifting when in the strain cycle a muscle spikes, timing can modulate force as much as rate in animals from cockroaches (56) to turkeys (57). For example, the same spike triplet can result in different force production depending on whether it occurs at onset of or during tetanus (58). Pressure production in bird respiratory muscle is sensitive to spike timing down to the millisecond scale. Across all these cases, the complex transformation of motor unit spike patterns into force gives potential for precise timing to convey rich information to control movement. Spike timing codes with corresponding timing sensitivity in muscle power production may be a prevalent feature in both individual motor programs and across species.

### Convergent mixed coding strategies for flight

An unexpected feature of the comprehensive motor program is the consistency in timing and count encoding across all the motor units (Fig. 4). Calcium imaging of the direct muscles controlling the wings in *Drosophila* showed evidence for two categories of muscle encoding: phasic muscles that are transiently active, especially during saccades, or tonic muscles that are continuously active (23). Flies may utilize a dichotomy of these exclusively phasic and tonic muscles organized into mixed functional groups, where at least one phasic and one tonic muscle is acting on each sclerite. In contrast, *Manduca sexta* utilizes muscles with a mix of spike timing and spike count encoding and usually have a larger, functionally dominant muscle (or muscles sharing innervation) in the group of muscles attached to sclerite as opposed to the similarly-sized muscles attached to each sclerite in flies (SI Appendix). Additionally, *Drosophila* fly at wing beat frequencies an order of magnitude higher than *Manduca sexta* and *Schistocerca nitens*. Larger size and longer wingbeat periods might allow for a single mixed timing and rate motor unit to have more power to control the sclerites. *Manduca sexta* also do not use saccades during flight, and muscles typically contract and relax on each cycle. While phasic and tonic calcium activation does not have the resolution of precise spiking activity, it does show a separation of timescales and the potential for separated mechanisms of coordination across muscles.

The information framework we use here is powerful in its generality, separates timing and count, and reveals the ubiquity, consistency, and coordination of spike timing. However, it does not indicate content of the signals on its own. Many different parameters in the motor signals could covary with torque and dissecting each component will require other approaches. We complement this information approach by examining specific patterns of spike count and spike timing related to torque in two example moths (Fig. S8-S9). DLMs varied with turn direction, but in a narrow timing window with low variance. This is consistent with their known control potential where changing individual spike times by as little as ± 4 ms can modulate the power output from 0% to 200% of normal and causally induce yaw torque (27). Overall left and right pairs of muscles shifted their timing differences across turns. Time separation in the DVM and modulating of the timing of the 3AX were also consistent with earlier work (41, 59). There were also some individual differences like in the basalar muscle where one moth increased the spike count for ipsilateral turns, while the other moth decreased the spike count. There may be significant individual variation in the particular control implementation each individual adopts even if the encoding strategy is conserved.

### Spike timing codes challenge motor circuit precision

Timing codes are limited by precision, both in the degree to which a spike can be reliably specified by the nervous system and reliably translated by the muscle and skeletal machinery into differential forces (4). The precise spiking of the indirect flight muscles has causal consequences for turning down to the sub-millisecond scale (27). We now understand that this likely extends across the entire motor program (Fig. 3*B*) and that coordination is achieved primarily thorough spike timing across muscles (Fig. 5*A-F*).

Given the relatively few spikes per wing stroke, spike count per period could easily be interpreted as a rate code in fast, periodically activated muscles like the hawk moth flight musculature, but there is a distinction between rate and spike count in some slow muscles with many spikes per cycle. In the slow cycle frequencies of the crustacean stomastogastric pyloric rhythm and stick insect strides, muscle force does not strictly follow rate encoding and depends on the specific number of spikes (19, 20). Timing codes are sometimes argued to be precise rate codes, but that would require drastic changes to spike rate in a very short time period for single spike codes, like the one present in the hawk moth DLM, and for codes that depend on specific spike patterns. For example, some slow muscles such as the radula closer in *Aplysia* show force dependence on specific patterns of spikes (60). Timing codes can be distinguished from rate or count codes by a specific pattern of spikes activated at a precise time in relation to a behavior (4). Using a phase timing code gave similar results to using an absolute timing code, due to there being little variation in the wing beat period, but it is also possible that information in phase and absolute timing may differ in systems where more variation exists in a characteristic movement period.

It is still unknown how peripheral temporal codes arise from higher brain areas, the central nervous system, or motor circuits in the spinal or ventral nerve cord. Precise timing could come from direct connections between sensory receptors and efferent units. In moths, there are rapid mechanosensory pathways from the antenna (61), wings (62), and potentially other organs that can provide reafference of movement that could be used for precision. In locusts, mechanical feedback from the tegula, a sensory organ depressed during each wing stroke, produces phase resetting in the flight motor pattern which coordinates the fore and hind wings (63). In flies, gap junctions exist between precise haltere mechanoreceptors (64) and steering muscles (65), producing very fast reflexes, which, in conjunction with fast feedback from wing mechanoreceptors, precisely patterns the activity of the first basalar muscle (66). However these reflexes are still influenced by visual commands that have to incorporate feedback passing through a number of central nervous system synapses (67). The millisecond scale resolution of the motor code poses a challenge even for neural processing that requires only a few synapses.

Precision may arise from central brain regions. Some pairs of bilateral muscles in *Drosophila* are innervated by motor neurons that receive input from the same circuitry in the nerve cord (68) which could give a proximal source of the left-right precision seen in *Manduca* downstroke muscles (27), but this alone is unlikely to be sufficient to account for the prevalence of timing codes. Central brain regions have been thought to encode information primarily by rate, but a cortical area for vocalization in song birds does show millisecond scale precision in encoding (25). Peripheral precision may also come from transforming a population code or remapping of dynamics distributed over large populations of neurons (32). Both the central nervous system and rapid peripheral sensorimotor pathways provide potential mechanisms for spike timing precision.

### Spike count does not inherently limit encoding

The prevalence of temporal coding in the moth motor program is not due to a limit in how much information can be encoded in spike count, since the spike count entropy was high enough to account for the total mutual information encoded by each muscle that spiked more than once per wing stroke (Fig. 5*H*). For the DVM and SA muscles, spike count would have to have no transmission error due to its entropy being similar in magnitude to the total MI, but for the 3AX and BA muscles, there could be transmission error and the spike count would still account for the total MI. Because much of the entropy in spike count is unused for encoding yaw torque, much of the variation in spike count must be ignored in the transformation from spiking activity to movement. The opposing trends in BA spike count from our two example moths may not affect the yaw torque (Fig. S8-S9). In fact, there is some evidence that muscle force is invariant to spike count in cockroach running (69).

While temporal codes are present both in faster, high frequency systems and slower, low frequency systems (18), count and rate codes are still used. Improved algorithms based on population rate codes for decoding motor implications of neural activity on a single-trial-basis have led to better neural prosthetic devices and brain-machine-interfaces (32, 70). Incorporating spike timing or pattern information shows promise for improving these devices by adding more information than what is present in just the rate code.

### Spike timing is essential to coordination

The moth motor program has redundancy in its information transmission. Yet, our estimate of the motor program information rate while accounting for shared information still enables the encoding of hundreds of unique states. Redundancy and synergy in information transmission have been explored in the sensory periphery and in central brain regions where there may be a trade-off between code efficiency and robustness to noise (71–73). Dimensionality reduction techniques are commonly used to study populations of neurons in motor brain areas or ensembles of muscles (31–33, 70, 74–76). The activation patterns of many muscles may be represented by low dimensional linear combinations of many muscle, "muscle synergies", that capture most of the variation (31, 33, 74, 75, 77). There is potential confusion of the terms synergy and redundancy, because muscle synergies are likely to share net redundant (not synergistic) information. In the moth, all combinations of muscles do share information (negative *II* values).

Analysis without considering timing may miss important structure in how brains coordinate movement. Previous investigations of muscle synergies could not assess coordination at the spike level, though modulation of msucle activation over longer timescales was an important component of synergies identified in frogs, cats, and humans (36, 74, 76). In the *Manduca* system nearly all of the coordination between muscles may be overlooked by not considering spike timing. All muscles are more coordinated in their timings than the DLMs that have zero entropy in spike count. The spike timings of the DLM muscles have previously been shown to exhibit a low degree of coordination in their code for yaw torque (33). This is consistent with our results, since we found that these two muscles have the least pairwise interaction information (Fig. 5*A-F*, Fig. S6). Not all information encoded by individual muscles was shared. In the moth motor program, each muscle has a small amount of independent motor information it can convey with count, while control encoded in timing is coordinated across multiple muscles (Fig. 5*B*).

The hawk moth motor program uses a precise, coordinated spike timing code along with a less informative but independent spike count code consistently in every muscle used to control the wings. Spike timing codes likely necessitate millisecond scale precision arising from either sensory feedback loops or central motor circuits. When combined with the growing number of specific examples of spike timing motor codes across vertebrates and invertebrates, the millisecond patterning of spikes can not be safely ignored or necessarily relegated to a few specific cases. Timing encoding in the most peripheral motor output may be more of a rule, not an exception.

## Materials and Methods

### Data Archival

The data used in this paper will be made available on Dryad (accession information upon publication).

### Electromyography (EMG) recordings from flight muscles

Moths (*Manduca sexta*) were obtained as pupae (University of Washington colony) and housed communally after eclosion with a 12-hour light-dark cycle. Naïve males and females (N = 7) were used in experiments conducted during the dark period of their cycle.We cold anesthetized moths before removing scales from the ventral and dorsal sides of their thoraxes. We made two small holes in the cuticle using insect pins and inserted two silver EMG wires to take differential recordings from the 10 indirect power muscles and direct steering muscles (Fig. S1). These 5 pairs of muscles comprise a nearly complete motor program for flight (SI Appendix). A common ground wire was placed in the abdomen. We imaged the external placement of silver EMG wires to ensure we targeted the correct muscles (Fig. S1). We also conducted post-mortem dissections on a subset of animals to verify wire placement. All images were captured with a Zeiss Stereo Discovery v.12 equipped with a Zeiss Axiocam 105 color camera.

### Experimental set-up

We tethered moths with cyanoacrylate glue to a 3D-printed ABS plastic rod rigidly attached to the force-torque (F/T) transducer (ATI Nano17Ti, FT20157; calibrated ranges: F_*x*_, F_*y*_ = ± 1.00 N; F_*z*_ = ± 1.80 N; *τ*_*x*_, *τ*_*y*_, *τ*_*z*_ = ± 6250 mN-mm). After tethering, we allowed 30 minutes for the moths to adapt to dark light conditions and recover from the surgery at room temperature before starting experimental recordings. We amplified the EMG signals using a 16-channel AC amplifier (AM Systems Inc., Model 3500) before acquisition with a NI USB-6259 DAQ board. We also acquired Gauge voltages from the F/T transducer a NI USB-6259 DAQ board. We sampled both the EMG and F/T transducer gauge voltages at 10000 Hz. We captured outputs from these DAQ boards using MATLAB (MathWorks).

## SI Methods

See the SI Methods for reporting on the visual stimulus used, spike train analysis, wing stroke alignment, and information theoretic estimates.

## ACKNOWLEDGMENTS

The authors thank Mark Willis, Tom Daniel, Ilya Nemenman, and Sam Sober for helpful discussions. This material is based upon work supported by the National Science Foundation Graduate Research Fellowship under Grant No. DGE-1650044 and Grant No. DGE-1444932. This work was also supported by an NSF CAREER (PoLS – 1554790) to S.S. and a Klingenstein-Simons Fellowship in the Neurosciences to S.S.

