## Supplementary Text for "Precise timing is ubiquitous, consistent and coordinated across a comprehensive, spike-resolved flight motor program"

### Timing is (almost) everything for coordinating a comprehensive, spike-resolved, flight motor program

**This PDF file includes:**

Supplementary text
Figs. S1 to S9
Tables S1 to S2
References for SI text citations

### Supporting Information Text

### SI Methods

**Visual stimulus.** We used an artificial robotic flower to provide visual stimulus to the moth during recording, similarly to previous studies where this stimulus was used to elicit smooth pursuit in hawk moths(1, 2). Unlike flies, moths do not use saccades to turn during flight. We actuated the flower flower in a purely horizontal, 1 Hz sinusoidal trajectory using precisely controlled servo motors (Phidgets, Inc.) connected to a 12 V DC power supply. We only considered trials where the moth was tracking the robotic flower. Different patterns of muscle activity have been observed for different types of behaviors, so we needed to control for tracking flight to ensure that we recorded and analyzed a consistent motor strategy(3, 4). To determine whether the moth was tracking the flower, we recorded high speed video at 250 fps above the moth (FASTEC IL4; 50 mm lens). We illuminated the working arena with an 850-nm IR light (Larson Electronics). We used black fabric and poster board to isolate the arena around the moth. We identified a tracking response based on the moth's head, wing, and abdominal motion in response to the flower's motion. For the trials where a visual tracking response was present, we computed the power spectral density of the yaw torque that the moth produced to determine whether a peak at 1 Hz was present, which was indicative of coherent motion with the flower. To ensure that this peak was not an artifact of the flower motion or other mechanical elements of our experimental set-up, we carefully isolated the F/T transducer from the robotic flower, speakers, and other vibrating machines in the experimental room.

**Spike train analysis.** We used Offline Sorter (OFS; Plexon) to detect the timing of spiking events in the EMG recordings. We first applied a threshold crossing method and then identified the peak in a short time window after threshold crossing. We manipulated the threshold value, waveform length, and inter-spike interval to maintain accurate and consistent detection. When multiple muscle signals were present on a single channel, we compared the raw signals from multiple channels. We referenced the literature considering typical shape and phase of each muscle signal (SI Appendix, Fig. S1) (3–7). When necessary, we high pass filtered data using a 4th order Butterworth filter (100 Hz cutoff). We combined all trials from the same individual for mutual information (MI) analysis.

**Wing stroke alignment.** The strain gauge voltages from the F/T transducer were transformed to calibrated forces and torques and translated to the point of the moth's attachment to the tether (the dorsal surface of the thorax). Spike timings during a wing stroke were referenced to the peak downward force in the z-direction during each wing stroke. The phase crossing was determined by filtering $F_z$ with an 8th order Type II Chebychev filter with a pass band of 3-35 Hz, which captures the natural wing beat frequency of *M. sexta* ( approximately 20 Hz in tethered flight). Both the torque and EMG data were segmented into wing strokes. For all following analyses, the raw yaw torque signal was low-pass filtered with a 4th order Butterworth filter (1000 Hz cutoff).

**Mutual information.** To reduce the dimensionality of the yaw torque in each wing stroke, we did a principal components analysis (PCA) on the torque waveforms within each individual. We also tested an alternative sampling method where we normalized the wing strokes to their period, and sampled yaw torque at several equally spaced phases during the wing stroke. Both methods give similar results. To determine the relative importance of count and timing encoding, we implemented a Kraskov $k$-nearest neighbors method of estimating MI previously used to analyze spikes from breathing muscles in songbirds (8–10). This method estimates the spike count MI before calculating additional spike timing MI (Eq. 1 in main text).

This method relies on the selection of an appropriate number of nearest neighbors ($k$) (8–10). We estimated the MI across different values of $k$. In most cases our estimates were insensitive to choice of $k$ (Fig. S2,S3), but in some cases, too small of a $k$ resulted in unstable estimates. We chose the smallest value of $k$ where estimates became stable in both k-space and for smaller subsets of data, $k = 4$. A few particular incidences required the full data, but the overall conclusions across muscles and moths were robust to data size. For all spike timing MI estimations, any spike count condition that occurred in less than $k$+1 wing strokes or fewer wing strokes than the dimensionality of $S_t$ were excluded from the summation.

We estimated error in our spike count MI estimates using the variance of $I(S_c, \tau)$ in non-overlapping fractions (for N = 1-10, data split into equal 1/N sets) of each individual moth's data set (10). To estimate error in spike timing MI estimates, similarly to above, we found the variance of each calculation $I(S_t, \tau|i)$. We then propagated the error through the weighted mean of $p(S_c = i)$. This method assumes no error in our estimation of the probability of each spike count condition. All the error estimates are at least an order of magnitude lower than the MI values, and are lower than the S.E.M. across individuals for all cases except the estimations of $I(S_r, \tau)$ for the DLMs, which approach $I = 0$ (Table S1).

**Pairwise MI and interaction information.** To investigate how MI is encoded across muscles, we estimated the joint mutual information between different pairwise combinations of muscles and the yaw torque response as the pairwise MI $I(S_A, S_B; \tau)$ (Eq. 2 in main text).

We estimated error in our pairwise spike count MI estimates using the variance of $I([S_{c,A}\ S_{c,B}], \tau)$ and the same methods as our individual muscle MI estimates. To estimate error in pairwise spike timing MI estimates, using the same data fractioning described above, we found the variance of each calculation of the second term of the pairwise MI equation (Eq. 2 in main text) and then propagated the error through the weighted mean of $p(i_A, i_B)$. All the error estimates are at least an order of magnitude lower than the pairwise MI values, and are lower than the S.E.M. across individuals (Table S2).

To compare the pairwise MI and individual muscle MIs, we used an interaction information measure $II$ (Eq. 3 in main text), which is the difference between the pairwise MI $I(S_A, S_B; \tau)$ (Eq. 2 in main text) and the sum of the individual muscle MIs (Eq. 1 in main text) for two muscles A and B (11).

We calculated this measure for separated spike count $II$ and spike timing $II$. The spike count interaction information is:

$$II_{count} = I([S_{A,c}\ S_{B,c}]; \tau) - (I(S_{A,c}; \tau) + I(S_{B,c}; \tau)) \quad [1]$$

This equation takes the spike count terms from both the pairwise MI estimate (Eq. 2 in main text) and the individual muscle MI estimates (Eq. 1 in main text). In the same way, the spike timing interaction information is:

$$\begin{aligned} II_{timing} = & \sum_{i_A=1}^{S_{A,c_{max}}} \sum_{i_B=1}^{S_{B,c_{max}}} p(i_A, i_B) I([S_{A,t}\ S_{B,t}]; \tau | (i_A, i_B)) \\ & - (\sum_{i_A=1}^{S_{A,c_{max}}} p(i_A) I(S_{A,t}; \tau | i_A) + \sum_{i_B=1}^{S_{B,c_{max}}} p(i_B) I(S_{B,t}; \tau | i_B)) \end{aligned} \quad [2]$$

Similarly to the full $II$, positive values of $II_{count}$ and $II_{timing}$ indicate net synergistic interactions between muscles A and B and negative values indicate net redundant interactions between muscles A and B.

**Spike count entropy and total motor program information.** We estimated the entropy of spike count using the direct method ((12, 13)):

$$H_c = -\sum_{i=1}^{S_{c,max}} p(S_c = i) log_2(p(S_c = i)) \quad [3]$$

The direct method estimates the entropy by the probability of each discrete state of the spike count condition $S_c = i$ up to the maximum number of spikes per wing stroke, $S_{c,max}$. The entropy is maximized by a uniform distribution, and minimized if only one spike count condition is present in the data.

To estimate the amount of information present in the motor program, we first calculated the sum of the total MI estimates of all muscles for each individual. This value does not account for redundancy, and therefore is an overestimation of the actual amount of information present in the motor program. We used three methods to determine a range of possible values for the total motor program MI. The minimum value of the total motor program MI, $MI_{min}$, was calculated assuming all interaction information values represented independent, non-overlapping shared information, so that the maximum possible amount of interaction information was subtracted from the sum of the individual muscle MIs. The maximum value of the total program MI, $MI_{max}$, was calculated assuming all interaction information values represented dependent, overlapping shared information, so only the highest $II(S_A, S_B; \tau)$ was subtracted from the sum of the individual muscle MIs:

$$MI_{max} = \sum_{A=1}^{10} I(S_A; \tau) - max(II(S_A, S_B; \tau) | A \neq B, B \in 1 - 10) \quad [4]$$

$A$ and $B$ represent each of the 10 muscles in the motor program. $I(S_A; \tau)$ is the total MI for each muscle $A$ (Eq. 1 in main text) and $II(S_A, S_B; \tau)$ is the $II$ for each possible combination of muscles (Eq. 2 in main text). To provide an intermediate, best estimate within this range, we reduced the sum of total MIs by the ratio of $II$ to $MI$ across all muscle pairs:

$$MI_{MP} = (1 + < \frac{II(S_A, S_B; \tau)}{I(S_A; \tau) + I(S_B; \tau)} >) \sum_{A=1}^{10} I(S_A \tau) \quad [5]$$

where $MI_{MP}$ is the final estimate for the total motor program MI. The maximum possible yaw torque entropy for each moth data set was determined by the number of wing strokes $w$ recorded for that individual:

$$H_{\tau,max} = log_2(w) \quad [6]$$

To estimate how precisely the motor program MI could define different yaw torque states, we used direct method estimations on the joint probability distribution of the yaw torque PCs for decreasing bin sizes. This was used to determine the entropy when the motor output was divided into increasing numbers of states. Once $H_{\tau,max}$ was reached, we did not estimate the entropy for smaller bin sizes. This gave a mapping between the number of yaw torque states and the yaw torque entropy. The motor program MI encodes information about yaw torque entropy, so this was used to estimate how many states of yaw torque can be differentiated or controlled by the spiking activity of the muscles under perfect transmission from spikes to yaw torque states.

### Identifying a Comprehensive Motor Program

**Overview of the Motor Program for Moth's Wings.** For this study, we focus on the synchronous muscles of the hawk moth *Manduca sexta*. We consider the primary muscles found in the mesothorax that are responsible for control and coordination of forewing movement in the hawk moth motor program. To identify these muscles, we first selected the primary wing-actuator muscles. Next, we identified the sclerites at the wing hinge that are well known to affect flight. We indentified the three muscles that are particularly important for moving these sclerites.

*Lepidoptera* in particular are known to control their wings using relatively few muscles compared to other insect orders (14). In contrast to *Drosophila* (> 200 Hz wing beat frequency), which change their wing position using two indirect, asynchronous muscles (neural stimulated once for many wing strokes) and approximately 17 direct, synchronous muscles (15), *Manduca* (20-25 Hz wing beat frequency) can adjust the firing patterns of all their flight muscles every wing stroke to change their wing position and control their flight. Before elaborating on the specific muscles that make up the motor program in detail, we will briefly summarize what is known about the mechanisms of hawk moth flight control to establish a basis for what is necessary to make up a comprehensive motor program for *Manduca sexta*.

Insects such as beetles, moths, flies and locusts control their flight by indirectly powering the gross deformation of wings through large strains of the thoracic cuticle. Insects then elicit fine-tuned changes to the wing stroke through the muscles acting on small sclerites at the wing base (e.g. pronation/supination and promotion/remotion of the wings) (6, 16). Hawk moths including *Manduca sexta* can adjust the position of their wings at rapid timescales from wing stroke to wing stroke (4, 17). Two indirect muscles on each side of these animals serve as primary initiators of wing upstroke and downstroke: the dorsolongitudinal muscle (DLM, downstroke) and the dorsoventral muscle (DVM, upstroke). Providing the major source of power for flight, these muscles make up the central drive of the hawk moth motor program. While these indirect muscles have sometimes been assumed to only function as motors for upstroke and downstroke, increasing evidence implicates the activation time of these muscles for turning (18–22). Sub-millisecond level changes in the activation time of the DLM have been shown to cause changes in the yaw torque (18), and phase differences in both the DLM and DVM have been implicated in control of yaw turns (19). Additionally, moths compensate for asymmetric wing damage by altering the phase difference between both DLM and DVM activation, indicating that moths use this time difference to adjust for the difference in roll torque produced by different sized wings (21). As a result, it is critical to record simultaneously muscles on both sides of the animal.

In addition to the indirect DLM and DVM, direct steering muscles have been implicated as important muscles for wing control. These muscles have the greatest influence on nuanced wing position because they pull on sclerites located at the wing hinge. The 2nd and 3rd axillary sclerites and the subalar sclerite have been recognized in the literature as important contributors to wing control in hawk moths (6, 7). The 3rd axillary sclerite is widely accepted as important for flight control in many insect genera. It articulates with the most posterior vein on the forewing, the anal vein. Contraction of the muscle that attaches to this sclerite primarily affects the remotion of the wing but may have other effects depending on the position of the wing in the hawk moth (7). Also associated with wing remotion, the 2nd axillary sclerite moves in concert with the subalar sclerite via a ligament attachment (7). In addition to its connection to the 2nd axillary sclerite, the subalar sclerite is loosely connected to both the fourth axillary sclerite and the posterior notal wing process (7). Due to its connections to other sclerites and its known capacity to limit the motion of the 3rd axillary sclerite, the subalar sclerite is particularly influential in the control of hawk moth wing motion (7). We included the upper 3rd axillary muscle (3AX) and the subalar muscle (SA) as components of our motor program because of their attachments to these two sclerites, which have important influences on wing position. The basalar sclerite and its associated muscles have been referred to as the primary antagonists of the 3rd axillary and subalar muscles, although an exact antagonist relationship likely oversimplifies the complex wing hing (5–7). The basalar sclerite is suspected to be involved in promotion, pronation, and depression of the wings in many different species of insects (6). Changes in the phase of activation of the primary basalar muscle (BA) is associated with turning maneuvers in multiple moth species, including *Manduca sexta* (4).Due to its relationship with the 3rd axillary and subalar muscles and the sclerite's capacity to affect wing position, we also included the primary basalar muscle in our motor program. For the remainder of this supplement, we consider each of the muscles in the mesothorax of *Manduca*. The default notation for muscles that we use in this document comes from Eaton 1988 (23). Our goal is to resolve notation issues across prior anatomical and electrophysiological references, to explain the inclusion or exclusion of muscles from our recordings, and to highlight which additional muscles are candidates for future investigations.

#### Indirect Muscles.

**Dorsolongitudinal Muscle (DLM)** We refer to muscle DL1 according to notation from Eaton 1988 when we refer to the DLM (23). This muscle spans the entire midsection of the thorax and is the most medial muscle. The five subunits run from the anterior scutum (or nodal process) posterior to the phragma. This muscle has been extensively studied in *Manduca* and other *Lepidoptera* (4, 18, 20, 24–26). The DLM is composed of five subunits each innervated by a single motor neuron. However, there has never been evidence of the individual units activating out of phase with one another (Fig. S1*A*), and the DLM recordings typically produce a single compound action potential. We follow previous convention, treating the whole DLM as a single motor unit and observing that DLM spike count remains constant at one spike per wing stroke (4, 6, 19–21, 25). More recently, sub-millisecond level time-shifts in the activation of the left and right DLM have been shown to cause changes in the power output of the muscle but the each individual muscle still behaves like a single motor unit (18). These findings suggest that the neuromuscular activity of the DLM is likely to play an important role in both generating wing depression and in more nuanced flight control. They also provide causal evidence that *Manduca sexta* alters the precise timing of muscle activation to control flight (4–7, 27).

Two other dorsolongitudinal muscles are also indirect muscles: DL2 and DL3 (we refer only to DL1 as the DLM, but DL2 and DL3 both fit into the dorsolongitudinal category anatomically)(23). Both are classified as indirect muscles because they do not insert on a wing sclerite. Muscle DL3 is a thin muscle that is found dorsal of DL1, originating near the median line and inserting on the postnotum. Its relatively small size compared to the DL1 means that it is unlikely to provide large strain deformation in the thoracic cuticle. DL2 (Dorsal Oblique according to Kammer's notation) is a muscle of comparable size to that of a single subunit of the DVM (see description below) (4). This muscle originates on the scutum and inserts on the phragma-like process of the postalar bridge (23). The Dorsal Oblique is not an elevator or depressor muscle. It may contribute to flight control by altering the mechanical properties of the exoskeleton. It may change the mechanical response to DLM and DVM activation, thereby indirectly affecting the wings. Kammer has noted changes in activation patterns in this muscle associated with turning maneuvers, which may indicate that this muscle plays a role in flight control (4). However, this muscle does not directly adjust wing placement and does not provide the major power for up-stroke and down-stroke, so any contribution to wing control is likely small in comparison to the indirect power sources or direct steering muscles. In asynchronous insects an analogous muscle, the pleurosternal, is thought to adjust the resonant frequency of the stretch activated flight motor (16, 28, 29), but in the hawk moth the flight program is neurogenic and wingbeat frequency follows the rhythmic excitation of the DL1. For these reasons, we choose to focus on only indirect muscles that contribute the primary mechanical power to the flight apparatus. We neglect DL3 and DL2 from our motor program although DL2 is likely to have an underappreciated role in synchronous flight control.

**Dorsoventral Muscle (DVM)** Found just lateral to the DLM are the indirect, upstroke, power muscles, the dorsoventral muscles. Several muscles have been referred to as elevator muscles in the literature. These muscles include DV1, DV2, DV3, DV4, and DV5 according to notation from Eaton 1988 (23). The elevator muscles occupy similar anatomical locations and are oriented such that they perform similar actions on the thorax. We chose to record a single subunit to represent the indirect upstroke muscles. Muscle DV1 according to Eaton's notation or the tergo-sternal muscle according to Kammer's notation is larger than the other elevator muscles (4). This muscle consists of three subunits, DV1abc, which are all innervated by the anterior branch of nerve IIN4. The anatomical location of DV1b makes it a relatively easy muscle to target ventrally for EMG recordings. Like the DL1 all three subunits of the DV1abc activate simultaneously with a compound action potential, although the separation of the individual subunits has not been as carefully examined electrophysiologically as in the DL1 (Fig S1*B*). As has been done in most prior recordings we treat the combined DV1 recording as the wing upstroke muscle signals. We target DV1b for our electrode insertion location and will always refer to this subunit when we refer to the DVM. Typical DVM recordings consist of one or two spikes per wing stroke, and may experience some slight shifts in timing relative to the DLM. This muscle consistently activates out-of-phase with the DLM.

Other muscle units that fall into the dorsoventral category include DV2, DV3, DV4, and DV5 according to Eaton's notation. We neglect DV3 (tergotrochanteral according to Kammer) because it extends down into the legs of the animal (23). We assume that any muscle extending to the legs is likely responsible for holding the legs close to the thorax to maintain proper flight posture and is not a major component of wing control at least in indirect fliers. DV2 originates on the scutum and inserts on the basicosta, while DV5 attaches to the scutum and inserts on the meron, respectively (23). Both Dv2 and Dv5 are oriented similarly to DV1. DV4 (another segment of the posterior tergocoxal muscle according to Kammer), found just lateral of DV2, DV3 and DV5, also originates on the scutum and inserts on the meron (the location of insertion of the subalar muscle). It is similar in size to DV2. For an in-depth analysis of the function of the dorsoventral muscles, it would be good to obtain recordings from DV2, DV4, and DV5 in addition to DV1 during flight behavior. DV2 and DV5 (anterior and posterior tergocoxal according to Kammer) are comparable in size to one unit of DV1. By recording from DV1b we capture the activity of a single subunit to represent the overall dorsoventral muscles' contributions to wing control. We chose on subunit that shares innervation with all the subunits in the largest indirect elevator muscle. The various units of the dorsoventral muscles are all referred to as elevators according to Kammer. We referenced Kondoh and Obara 1982 when translating between Eaton and Kammer's notation(30).

### Direct Muscles.

**Pleurodorsal Muscles: 3rd Axillary** The 3rd axillary muscle (3AX), corresponding to muscle PD2 according to Eaton's notation, inserts on the 3rd axillary sclerite and is known to fluctuate its activity patterns during turning maneuvers (23, 31, 32). Kammer splits this muscle into three subunits: the upper, middle and lower 3rd axillary, each of which have different angles of origin. The segment of the muscle, which Eaton considers PD2ab comprises the upper 3rd axillary, and Kammer splits Eaton's PD2c into two separate subunits: the middle and lower 3rd axillary muscles (7, 23). We use notation from Rheuben and Kammer 1987 for this muscle (7). The upper and lower 3rd axillary muscles consist of intermediate fibers, while the middle 3rd axillary muscle consists of tonic fibers. The activity differs between the three subunits (7, 14, 33). The upper and lower 3rd axillary muscles exhibit large amplitude spiking activity during flight. They are innervated by fast axons projecting from the same nerve branch (IIN5c). The tonic muscle, the middle 3rd axillary, is innervated by a slow axon (IIN2a). There is evidence of muscle activity both during flight and at rest (7, 32). The amplitudes of activations of the middle 3rd axillary are much smaller than those of the upper and lower 3rd axillary muscle, and are therefore quite difficult to record (32). We chose a single subunit to represent all muscle activity responsible for moving the 3rd axillary sclerite. Following this logic and prior convention (4, 5), we record from the largest intermediate fiber subunit, the upper 3rd axillary, which attaches on the anepisternum (Fig. S1*C*).

It should be noted that extensive work has previously investigated the role of various 3rd axillary subunits in both the meso- and meta-thoracic segments during flight (4, 5, 7, 32, 34). Rheuben and Kammer note that because of the differing angles of origin, the three subunits are likely to have differing affects on the motion of the 3rd axillary sclerite. Additionally, the metathoracic 3rd axillary muscle is known to change the position of the hind-wings. A good extension to our study of the 3rd axillary muscle would be to record from the lower 3rd axillary in both the meso- and the meta- thoracic 3rd axillary subunits to determine whether activity in these muscles encodes additional information about torque output. We focus on the mesothoracic segment for our motor program because the forewings (the largest wings) attach to that segment of the thorax.

**Pleurodorsal Muscles** The other pleurodorsal muscles noted by Eaton include PD1, PD3, PD4, and PD5 (23). PD1, PD4, and PD5 are very small in size and their influence on wing control is likely negligible. PD3 inserts on the 1st axillary sclerite and is comparable in size to one of the subunits of the 3rd axillary muscle. Kammer mentions that this muscle may play a role in wing control for a saturniid moth, *Rothschidia jacobeae*, but claims that this muscle does not exist in hawk moths (5). In contrast, Eaton's documentation on the muscles supports the presence of this muscle in *Manduca sexta*. However, he provides no information about innervation to this muscle (23). According to Wendler, PD3 deteriorates within 48 hours after moth ecloses (32). For these reasons, we assume this muscle does not play a role in flight control. Moreover Kammer indicates that PD3 in saturniid moths may consist of tonic muscle fibers like those of the middle 3rd axillary muscle in hawk moths. Because the role of PD3 in wing control has not been well established in the literature, and due to its suspected fiber-type in saturniid moths, we neglect the role of this muscle in the motor program.

**Pleuroventral Muscles: Basalar Muscle** Three pleuroventral muscles, PV1, PV2, and PV3, come together to make up different parts of the muscle that Kammer refers to as the basalar muscle. The basalar is thought to play a role in wing supination and to work with the subalar muscle to provide some power for downstroke (4). This muscle has been shown to exhibit substantial fluctuation in its phase of activation relative to the DLM and in the number of activation signals per wing stroke (Fig. S1*D*). The phase of activation has been correlated to wing position and turning behavior in *Manduca* and other genera, indicating that this muscle's activity likely encodes the animal's motor output (4, 5). When we refer to the BA from our motor program, we refer to PV2, or the coxobasalar muscle. Muscle PV1 is flat and short, PV2 originates on the basicosta and inserts on the basalar tendon cap, and PV3 extends deep into the legs (23). We neglect PV1 because of its small size relative to PV2, and we neglect PV3 because it extends into the legs. Kammer finds no evidence of variation in activity between the three basalar subunits. By recording from PV2, we record a representation of overall activity of the basalar muscle for wing control (4).

**Pleuroventral Muscles: Subalar Muscle** The muscle units that Kammer refers to as the subalar muscle consist of muscles PV4 and PV5. According to Eaton, both muscles attach at the meron; PV4 inserts on the subalar sclerite, and PV5 inserts on the subalar tendon cap (located close to the sclerite) (23, 24). Kammer does not mention a distinction between these two muscles (4, 17). Details further distinguishing these two muscles are not well documented in the literature. We use Kammer's notation, referring to both PV4 and PV5 when we refer to the SA. We record from PV4 and PV5 as a single subalar muscle subunit (Fig. S1*E*). The activity of the subalar muscle has been attributed to wing remotion and promotion as well as direct wing depression (4). Changes in the activation timing of the SA relative to the DLM are associated with wing remotion and promotion (4). This muscle typically fires nearly in-phase with the DLM, although our recordings show a range of timing variation in the SA, which is comparable with that of the DLM. The muscle consists of phasic fiber type and is innervated by the same nerve as the upper and lower 3rd axillary, which projects fast axons to the subalar muscle (7, 33). The subalar sclerite connects to the 2nd axillary sclerite via two ligaments. Kammer notes the significant role that the 2nd axillary sclerite plays in wing control, but we have not found documentation of any muscles inserting on this sclerite. The physical connection between the subalar and 2nd axillary sclerites further supports that the subalar muscles play an important role in wing control.

**Pleuroventral Muscles** Other pleuroventral muscles include PV6 and PV7. PV6 is located towards the anterior ventral part of the mesothorax. It does not attach or insert near the winghinge (23). For this reason, we neglect this muscle in our motor program. PV7 originates at the pleural ridge, near the pleural wing process, and it inserts on the furca via a long tendon (23). The pleural ridge and the pleural wing process come together as a fulcrum for the wing. We neglect PV7 from our motor program because of lack of prior recording and because sclerites located around the wing fulcrum are better positioned and more likely to contribute to wing control than the fulcrum itself.

**Pleural Muscles** The two pleural muscles include P1 and P2. Muscle P1 is a short muscle that inserts on the subalar tendon cap in the mesothoracic segment after originating on the furca in the metathoracic segment. P2 is a small muscle that attaches posteriorly to P1 to the furca from the mesothoracic segment. It inserts on the pleural ridge on the dorsal side of the metathoracic segment (23). We neglect these muscles in our motor program because these two muscles are not solely mesothoracic wing muscles- they connect the meso- and meta- thoracic segments (23)

**Sternocoxal Muscles** Two of the four sternocoxal muscles (SC2 and SC3) are neglected because they are found deep in the legs. We assume muscles extending deep into the legs serve to hold legs in position during flight. SC1 is a short muscle that is found ventral to the wing hinge. We neglect this muscle because its distance from the wing hinge makes it unlikely to play a significant role in wing control. SC4 is a very thin muscle that originates on the furca and inserts on the meron. Because of the small diameter of this muscle (muscle force production is known to correlate with the diameter of the muscle) and its distance from the wing hinge, we neglect SC4 from our motor program (23).

**Coxal Muscles** Coxal muscles include CXM1, CXM2, and CXM3. Each of these muscles are found in the leg (23). We neglect all leg muscles from our motor program, so we do not consider these.

**Intersegmental Muscles** The intersegmental muscle, IS, is a very small muscle that is found at the posterior end of the mesothoracic segment. This muscle's size and its distance from the wing hinge make it unlikely to significantly contribute to wing control (23).

**Ventrolongitudinal Muscles** Both the ventrolongitudinal muscles, IIVL1 and IIVL2 are found in the ventral most region of the mesothoracic segment. We neglect these two muscles from our motor program because their distance from the wing hinge makes them unlikely to play an important role in wing control(23).

**Summary.** We have reviewed all of the muscles found in the mesothoracic segment of *Manduce sexta.* We identified five bilateral pairs of muscles to represent moth's complete motor program. We record from the DLM and DVM - the indirect muscles that best known for their roles wing control and the 3AX, BA, and SA - the direct muscles that are best known for their roles in wing control. Additionally, we provided a rationale for all muscles that are we did not record in the comprehensive motor program, although we highlight a couple of muscles that are the most likely to provide any missing information in the wing motor program, notably the dorsal oblique and the lower 3rd axillary.

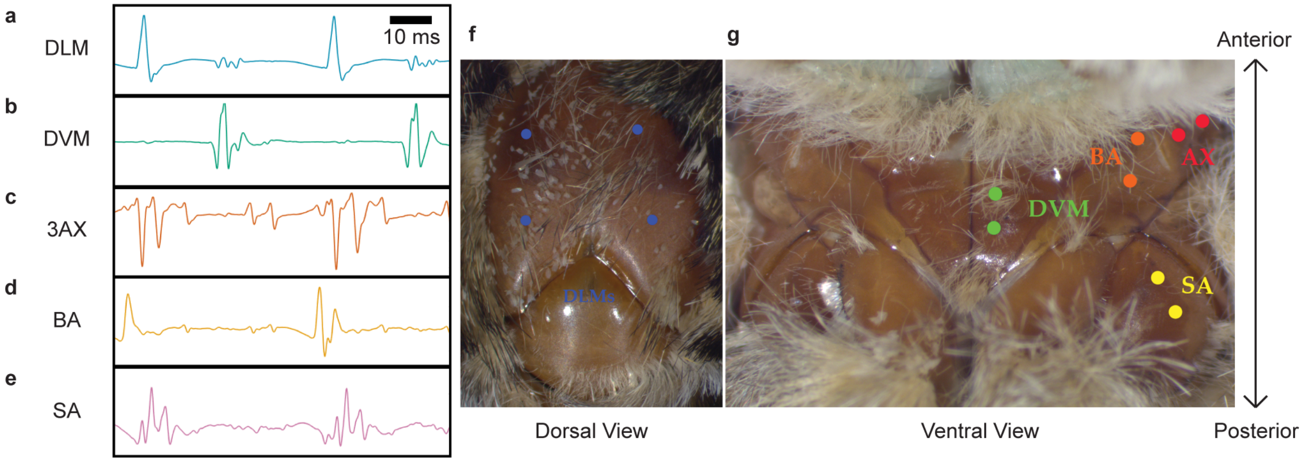


**Fig. S1.** Example recordings of two wing strokes from one of each of the the five bilateral muscle pairs in our motor program and wire placements. (*A*) The dorsolongitudinal muscle (DLM); (*B*) the dorsoventral muscle (DVM); (*C*) the third axillary muscle (3AX); (*D*) the basalar muscle (BA); (*E*) and the subalar muscle (SA) are all shown. All muscles had two silver EMG wires inserted through the cuticle of the thorax in order to obtain differential recordings of muscle activity. A ground wire was placed in the abdomen of the moth. (*F*) Dorsal view of the moth thorax showing DLM (blue) wire placement for the left and right sides in blue. (*G*) Ventral view of the moth thorax showing DVM (green), BA (orange), 3AX (red), and SA (yellow) wire placement for the left side. Images are aligned from top to bottom on the anterior-posterior axis. Wire placement in both panels was determined by overlaying wire insertion points in later images onto the thorax images shown, which were taken before surgery.

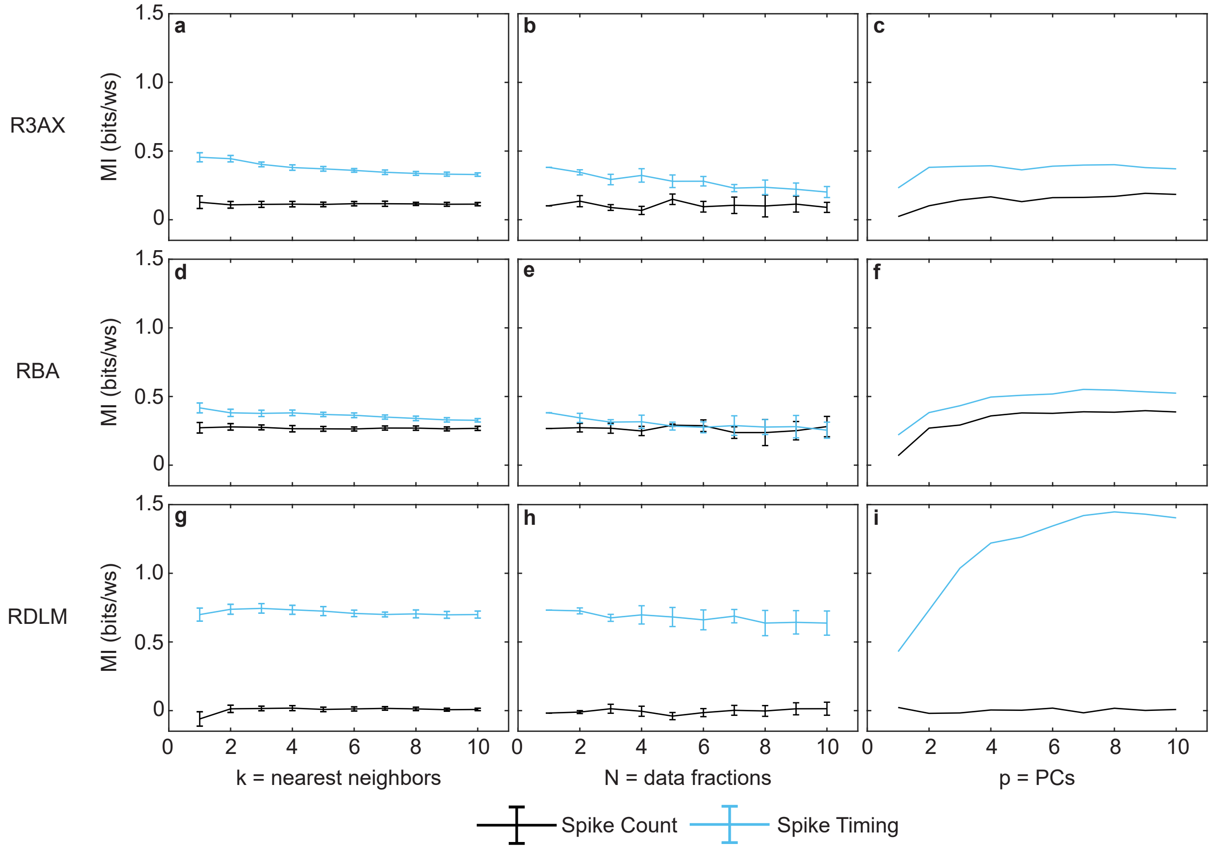


**Fig. S2.** Examples from one moth of MI estimates at different choices of $k$, the number of nearest neighbors; $N$, the number of data fractions; and $p$, the number of PCs included. We report the spike count (in black) and spike timing (in blue) MI estimates for the R3AX (*A-C*), RBA (*D-F*), and RDLM (*G-I*) muscles for $k$ (*A,D,G*), $N$ (*B,E,H*), and $p$ (*C,F,I*). 3AX and BA have the highest dimensionality in timing (high spike counts in some wing strokes) of all the muscles, so these are likely the most difficult in which to obtain stable estimates. The DLM has the lowest dimensionality in timing. For the estimates reported in the main text, we chose $k$ = 4 nearest neighbors due to its relative stability across the data sets.

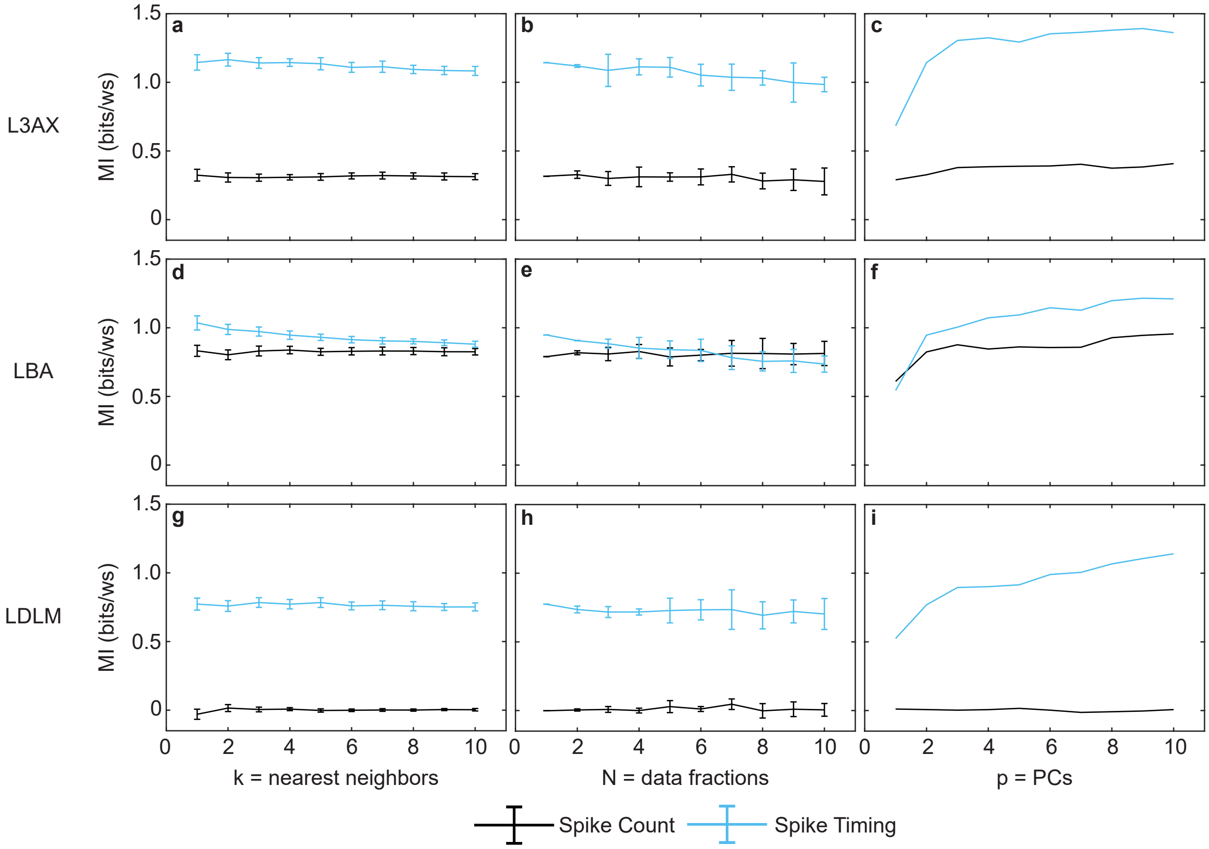


**Fig. S3.** Examples from one moth of MI estimates at different choices of $k$, the number of nearest neighbors; $N$, the number of data fractions; and $p$, the number of PCs included. We report the spike count (in black) and spike timing (in blue) MI estimates for the L3AX (*A-C*), LBA (*D-F*), and LDLM (*G-I*) muscles for $k$ (*A,D,G*), $N$ (*B,E,H*), and $p$ (*C,F,I*).

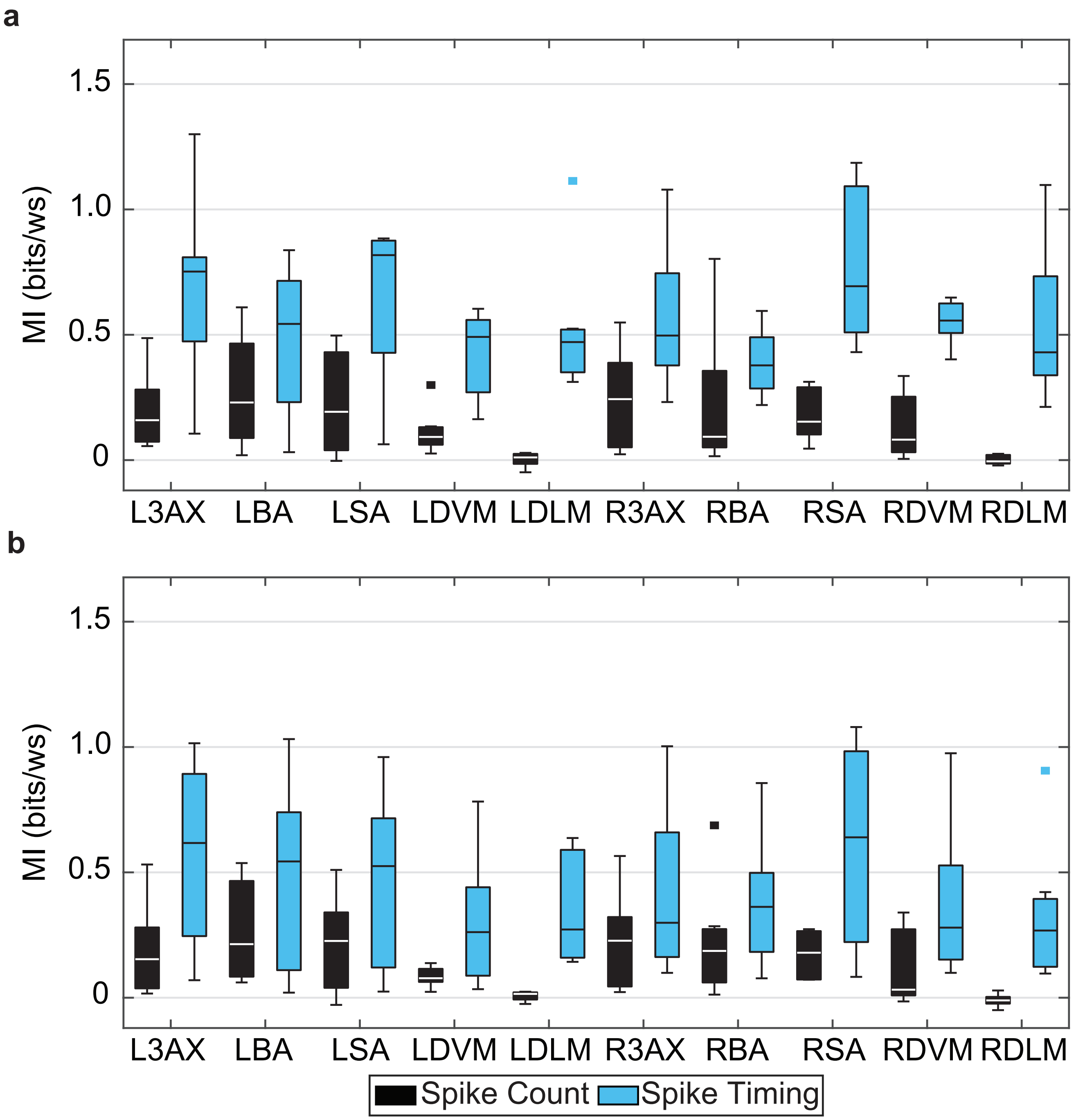


**Fig. S4.** (*A*) MI estimates for spike count (black) and spike timing (blue) with yaw torque represented using scores of just PC1 across individuals (N = 7). Box plots report the median as the center line in the box, which marks the 25th and 75th percentiles. Whiskers are range of all points that are not considered outliers (square points). Spike count MI is less than spike timing MI (two-way ANOVA comparing timing vs. count for all muscles except DLM: count vs. timing, $p < 10^{-}12$; muscle ID, p = 0.24; interaction, p = 0.19). Spike timing MI is significantly greater than spike count MI in most paired comparisons within muscles (paired t-tests: $p < 0.02$ for all muscles except the LBA, p = 0.16, and RBA, p = 0.17. Wilcoxon signed rank tests: $p < 0.04$ for all muscles except the LBA, p = 0.22, and RBA, p = 0.22). (*B*) MI estimates for spike count (black) and spike timing (blue) with yaw torque represented using the wing-stroke averaged yaw torque, rather than the PC scores, across individuals (N = 7). Box plots report the median as the center line in the box, which marks the 25th and 75th percentiles. Whiskers are range of all points that are not considered outliers (square points). Spike count MI is less than spike timing MI (two-way ANOVA comparing timing vs. count for all muscles except DLM: count vs. timing, $p < 10^{-}8$; muscle ID, p = 0.55; interaction, p = 0.89). Spike timing MI is significantly greater than spike count MI in most paired comparisons within muscles (paired t-tests: $p < 0.05$ for all muscles except the LBA, p = 0.15, and RBA, p = 0.18. Wilcoxon signed rank tests: $p < 0.04$ for all muscles except the LBA, p = 0.22, and R3AX, p = 0.08, and RBA, p = 0.22).

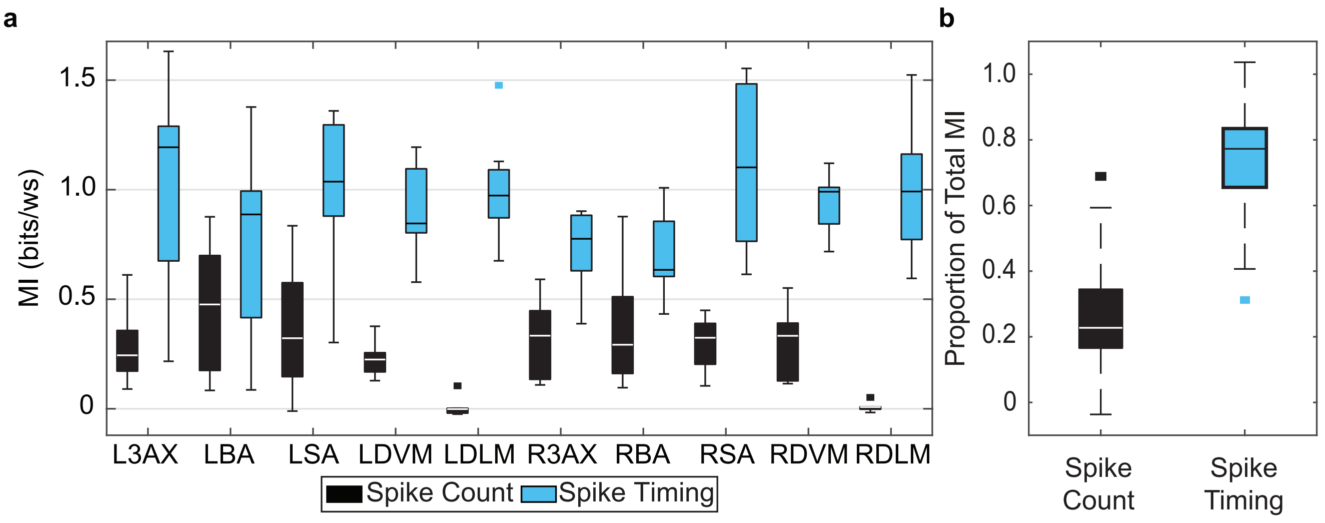


**Fig. S5.** (*A*) MI estimates for spike count (black) and spike timing (blue) with yaw torque represented using scores of the first 3 PCs across individuals (N = 7). Box plots report the data as previously described in Fig. S4. Whiskers are range of all points that are not considered outliers (square points). Spike count MI is less than spike timing MI (two-way ANOVA comparing timing vs. count for all muscles except DLM: count vs. timing, $p < 10^{-17}$; muscle ID, p = 0.78; interaction, p = 0.25). Spike timing MI is significantly greater than spike count MI in most paired comparisons within muscles (paired t-tests: $p < 0.03$ for all muscles except the LBA, p = 0.11. Wilcoxon signed rank tests: $p < 0.05$ for all muscles except the LBA, p = 0.11). (*B*) The proportion of spike count MI (black) and spike timing MI (blue) to total MI, respectively, across 8 muscles (DLM excluded) and 7 individuals. No significant difference was found in the magnitude of spike count MI of all muscles excluding the DLM (one-way ANOVA: p = 0.68; Kruskal-Wallis test: p = 0.95) or spike timing MI of all muscles (one-way ANOVA: p = 0.50; Kruskal-Wallis test: p = 0.38). No significant difference was found in the proportion of spike timing MI to total MI in all muscles excluding the DLM (one-way ANOVA: p = 0.17; Kruskal-Wallis test: p = 0.50).

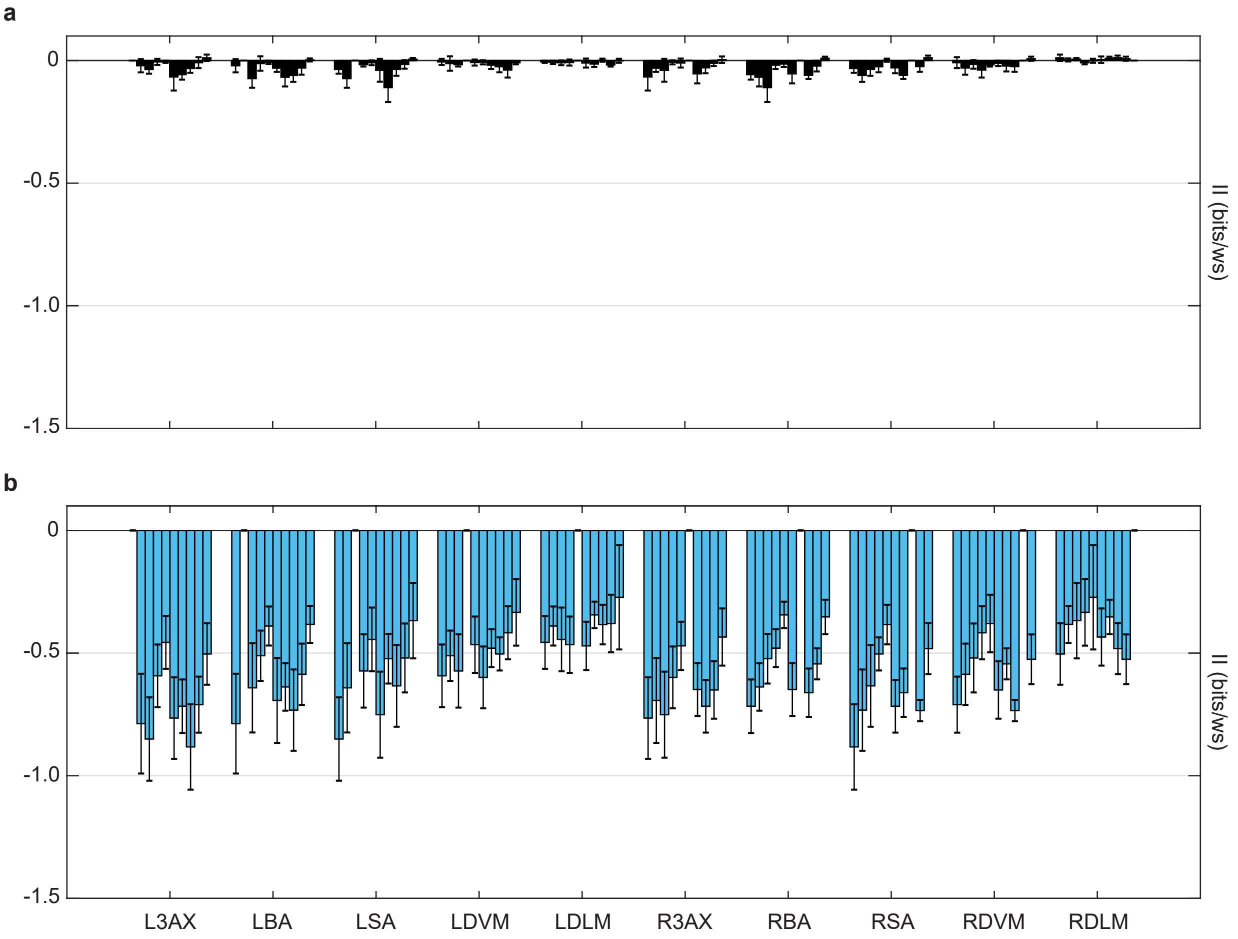


**Fig. S6.** Spike count and spike timing interaction information ($II$) estimates for pairwise combinations of muscles. Mean $\pm$ S.E.M. of (*A*) spike count $II$ (in black) and (*B*) spike timing $II$ (in blue) across N = 7 moths. This is the same data as in Fig. 4*B-C* in the main text, but with each row broken out to show the S.E.M. across individuals. Each group of bars is ordered to show pairwise interactions between the muscle labeled on the x-axis and, in order, the L3AX, LBA, LSA, LDVM, LDLM, R3AX, RBA, RSA, RDVM, and RDLM muscles. The identity interaction was not calculated for a muscle with itself.

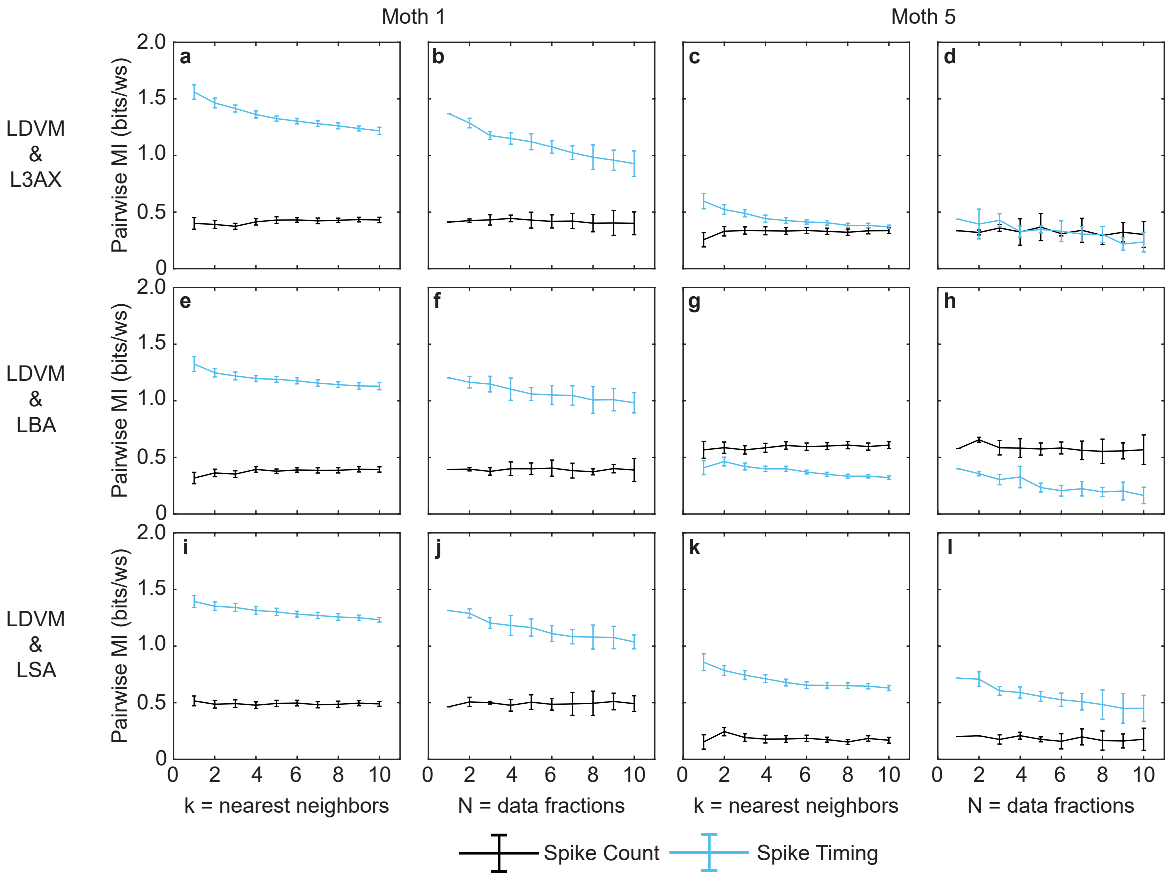


**Fig. S7.** Examples from two moths of pairwise MI estimates at different choices of $k$, the number of nearest neighbors; and $N$, the number of data fractions. For each moth (Moth 1: *A-B,E-F,I-J* and Moth 5: *C-D,G-H,K-L*), we report the spike count (in black) and spike timing (in blue) pairwise MI estimates for the RDVM and R3AX (*A-D*), RDVM and RBA (*E-H*), and RDVM and RSA (*I-L*) muscles for $k$ = 1-10 and $N$ = 1-10. For the estimates that were used to estimate interaction information reported in the main text, we chose k = 4 nearest neighbors due to its relative stability across the data sets.

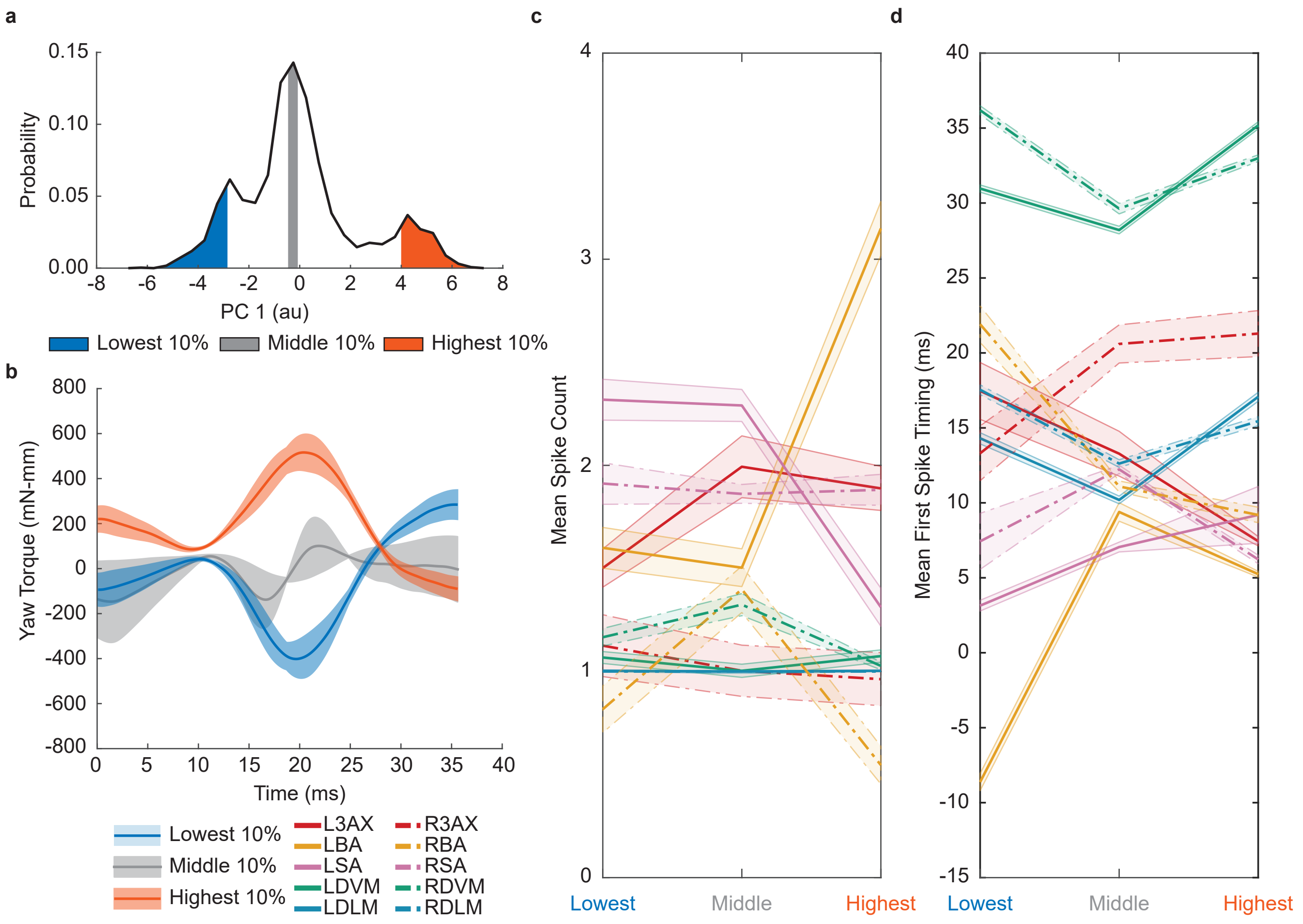


**Fig. S8.** Reconstructions using the first 2 PCs of yaw torque correlates with spiking activity in the same moth as in main text Fig. 2. (*A*) This is the same data as presented in main text Fig. 2*A*. Histogram of the scores of PC1. The lowest decile (0-10%), the middle decile (45-55%), and highest decile (90-100%) are shaded in blue, grey, and orange, respectively. (*B*) Mean $\pm$ S.D. of the reconstructions of wing strokes from the lowest (blue), middle (grey), and highest (orange) deciles using the data set mean and the projections of scores onto the first 2 PCs. This differs from the main text Fig. 2*B* because it does not display the raw torque waveforms, instead showing variation in the reconstructions. The lowest decile of PC1 corresponds to the left yaw turns, the middle decile corresponds to straight flight, and the highest decile corresponds to right turns. Mean $\pm$ 95% C.I. spike counts (*C*) and the first spike timing (*D*) in the lowest, middle, and highest deciles for all muscles. In this moth, the spike count of the basalar muscle and the spike count of the subalar muscle appear to change from the lowest to highest torque deciles. The ipsilateral subalar muscle has a higher mean spike count during turning. The spike count of the ipsilateral basalar muscle seems to decrease for turns, which is a trend that is consistent with Kammer 1971 (4). For the mean time of the first spike, the ipsilateral DLM, DVM, and SA spikes precede the contralateral spikes in the turning deciles. This trend is consistent with our previous results from stimulating the DLM, where we found that stimulating a spike in the left DLM before the time that the right DLM spike naturally occurs causes the moth to turn left and vice versa (18).

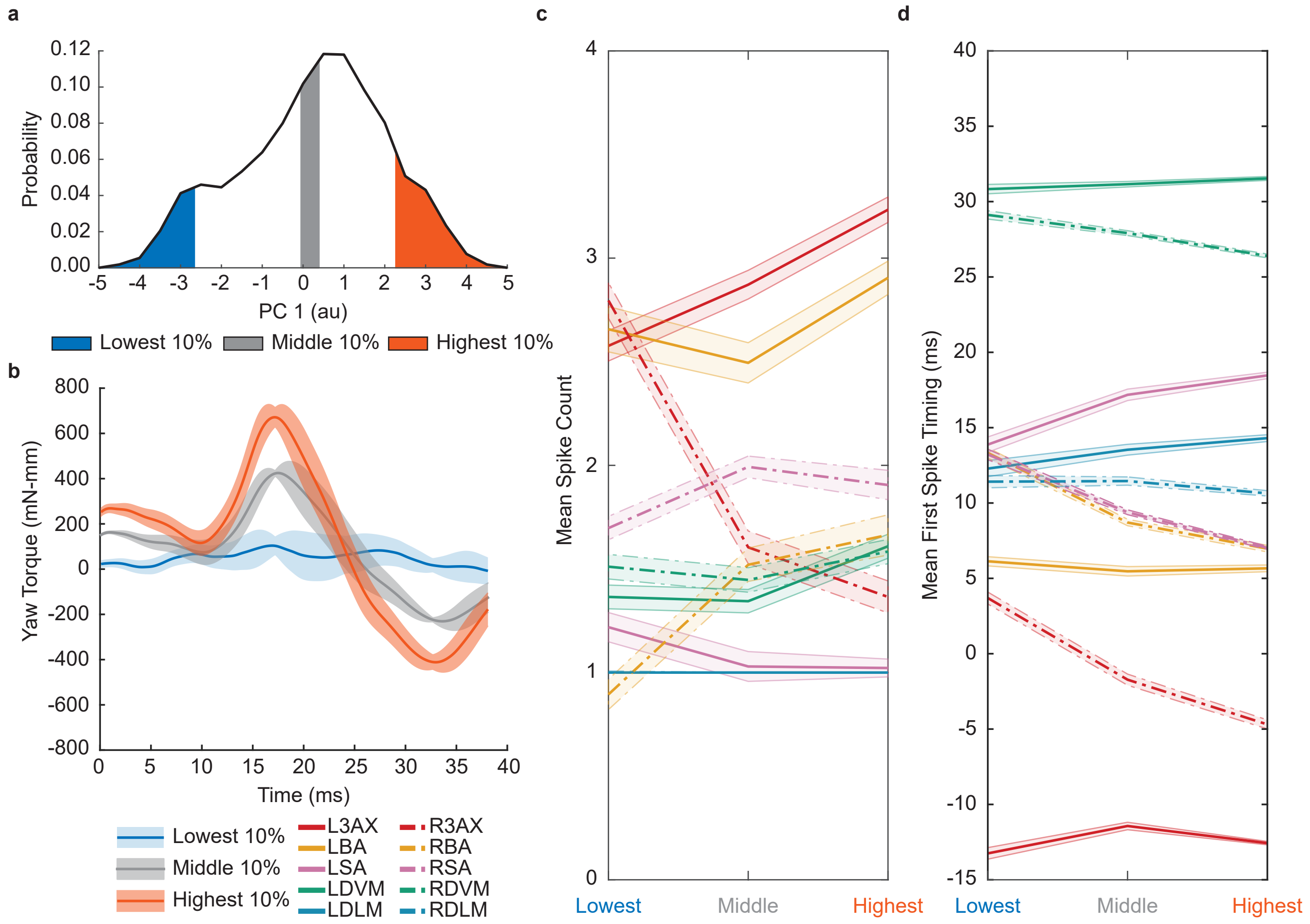


**Fig. S9.** Reconstructions using the first 2 PCs of yaw torque correlates with spiking activity in a different moth than in main text Fig. 2. (*A*) Histogram of the scores of PC1. The lowest decile (0-10%), the middle decile (45-55%), and highest decile (90-100%) are shaded in blue, grey, and orange, respectively. (*B*) Mean $\pm$ S.D. of the reconstructions of wing strokes from the lowest (blue), middle (grey), and highest (orange) deciles using the data set mean and the projections of scores onto the first 2 PCs. The lowest decile of PC1 corresponds to the straight flight, the middle decile corresponds to middle right turns, and the highest decile corresponds to the hardest right turns. Mean $\pm$ 95% C.I. spike counts (*C*) and the first spike timing (*D*) in the lowest, middle, and highest deciles for all muscles. In this moth, the mean spike count of the right basalar muscle increases from the lowest to highest torque deciles. Interestingly, in contrast to the moth from Fig. S8, this trend opposes Kammer's note that for the hardest turns, the ipsilateral BA did not spike (4). Similarly to the moth from Fig. S8, for the mean time of the first spike, the ipsilateral DLM, DVM, and SA spikes precede the contralateral spikes in the turning deciles. The timing difference is increased for the hardest right turns. Again, this trend is consistent with our previous results from stimulating the DLM, where we found that stimulating a spike in the left DLM before the time that the right DLM spike naturally occurs causes the moth to turn left and vice versa (18).

**Table S1. Error estimates for individual MI estimates for each muscle (Eq. 1 in the Main Text). Values were averaged across N = 7 moths.**

| Muscle | Count MI (bits/ws) | Timing MI (bits/ws) |
|---|---|---|
| L3AX | 0.0266 | 0.0309 |
| LBA | 0.0260 | 0.0237 |
| LSA | 0.0227 | 0.0295 |
| LDVM | 0.0217 | 0.0329 |
| LDLM | 0.0199 | 0.0344 |
| R3AX | 0.0279 | 0.0293 |
| RBA | 0.0246 | 0.0282 |
| RSA | 0.0258 | 0.0318 |
| RDVM | 0.0248 | 0.0331 |
| RDLM | 0.0206 | 0.0323 |

**Table S2. Error estimates for pairwise spike count MI (upper triangle in black, bits/ws) and pairwise spike timing MI (lower triangle in blue, bits/ws) estimates for each muscle (Eq. 2 in the Main Text). Values were averaged across all moths (N = 7) for each pairwise interaction.**

| - | L3AX | LBA | LSA | LDVM | LDLM | R3AX | RBA | RSA | RDVM | RDLM |
|---|---|---|---|---|---|---|---|---|---|---|
| L3AX | - | 0.0322 | 0.0276 | 0.0316 | 0.0292 | 0.0308 | 0.0320 | 0.0296 | 0.0289 | 0.0282 |
| LBA | 0.0246 | - | 0.0329 | 0.0326 | 0.0282 | 0.0311 | 0.0316 | 0.0320 | 0.0304 | 0.0283 |
| LSA | 0.0271 | 0.0242 | - | 0.0294 | 0.0267 | 0.0293 | 0.0306 | 0.0313 | 0.0317 | 0.0263 |
| LDVM | 0.0283 | 0.0268 | 0.0290 | - | 0.0259 | 0.0301 | 0.0288 | 0.0310 | 0.0271 | 0.0283 |
| LDLM | 0.0312 | 0.0308 | 0.0317 | 0.0313 | - | 0.0291 | 0.0292 | 0.0274 | 0.0246 | 0.0217 |
| R3AX | 0.0264 | 0.0230 | 0.0254 | 0.0285 | 0.0296 | - | 0.0294 | 0.0304 | 0.0314 | 0.0284 |
| RBA | 0.0252 | 0.0235 | 0.0290 | 0.0278 | 0.0315 | 0.0231 | - | 0.0310 | 0.0298 | 0.0290 |
| RSA | 0.0271 | 0.0247 | 0.0288 | 0.0306 | 0.0316 | 0.0264 | 0.0257 | - | 0.0298 | 0.0287 |
| RDVM | 0.0286 | 0.0276 | 0.0298 | 0.0309 | 0.0338 | 0.0272 | 0.0280 | 0.0284 | - | 0.0263 |
| RDLM | 0.0310 | 0.0313 | 0.0315 | 0.0347 | 0.0336 | 0.0309 | 0.0306 | 0.0308 | 0.0350 | - |
